# Protease secreting *Nannochloropsis limnetica* can valorise invasive plants through circular water-use and accumulate neutral lipids

**DOI:** 10.1101/2025.11.29.691304

**Authors:** A. Iyer, P.G. Heneghan, E.T. Dillion, R. Halim

**Affiliations:** Conway Institute of Biomolecular Research, University College Dublin, Belfield, Ireland; University College Dublin, School of Biosystems and Food Engineering, Belfield, Ireland

**Keywords:** Nannochloropsis, protease, invasive plants, wastewater, mixotrophic

## Abstract

*Nannochloropsis limnetica* can be cultivated on a wide range of effluents and generate high value lipids. Using a fed-batch system, this microalgae was used to demonstrate circular reuse of water to valorise protein depleted fractions (PDF) generated from Gorse leaf protein concentrate (LPC) production. Protein, phenolics and sugar content in the PDF was measured over seven days and corresponding microalgal lipid content was profiled. Extracellular protease activity was also screened to understand how *N. limnetica* digests and utilises proteins in the medium. A signal peptide-based approach predicted seven proteases expected to be secreted.

*N. limnetica* could rapidly deplete nutrients (k_sugar_=0.20 ± 0.06 day^−1^ and k_proteins_=0.17 ± 0.04 day^−1^) from the wastewater which could be reused for subsequent LPC extraction processes without any impact on recovery or purity. Phenolics levels remained unaffected during microalgal growth and accumulated in the water across each extraction cycle. Microalgal neutral lipid content (∼132.8 mg_lipid_ g^−1^ ) was higher in the effluent compared to standard BBM (∼114.5 mg_lipid_ g^−1^ ). Casein agar plates and 1D zymogram confirmed extracellular protease production by *N. limnetica*, although their identities remains unclear owing to conflicting / fragmented data from the LC/MS analysis.

This work shows that *N. limnetica* is able to secrete proteases to digest and utilise residual proteins in the medium, and can be employed to valorise agricultural wastewater. Identification of the proteases will require further investigation.

**Highlights:**

1. *N. limnetica* can rapidly deplete residual nutrients from LPC effluents.
2. *N. limnetica* accumulates higher neutral lipids with LPC effluents versus BBM.
3. *N. limnetica* secretes extracellular proteases to digest proteins in the medium.
4. Water can be circularly reused between LPC production and *N. limnetica* cultivation.

**Graphical abstract:** 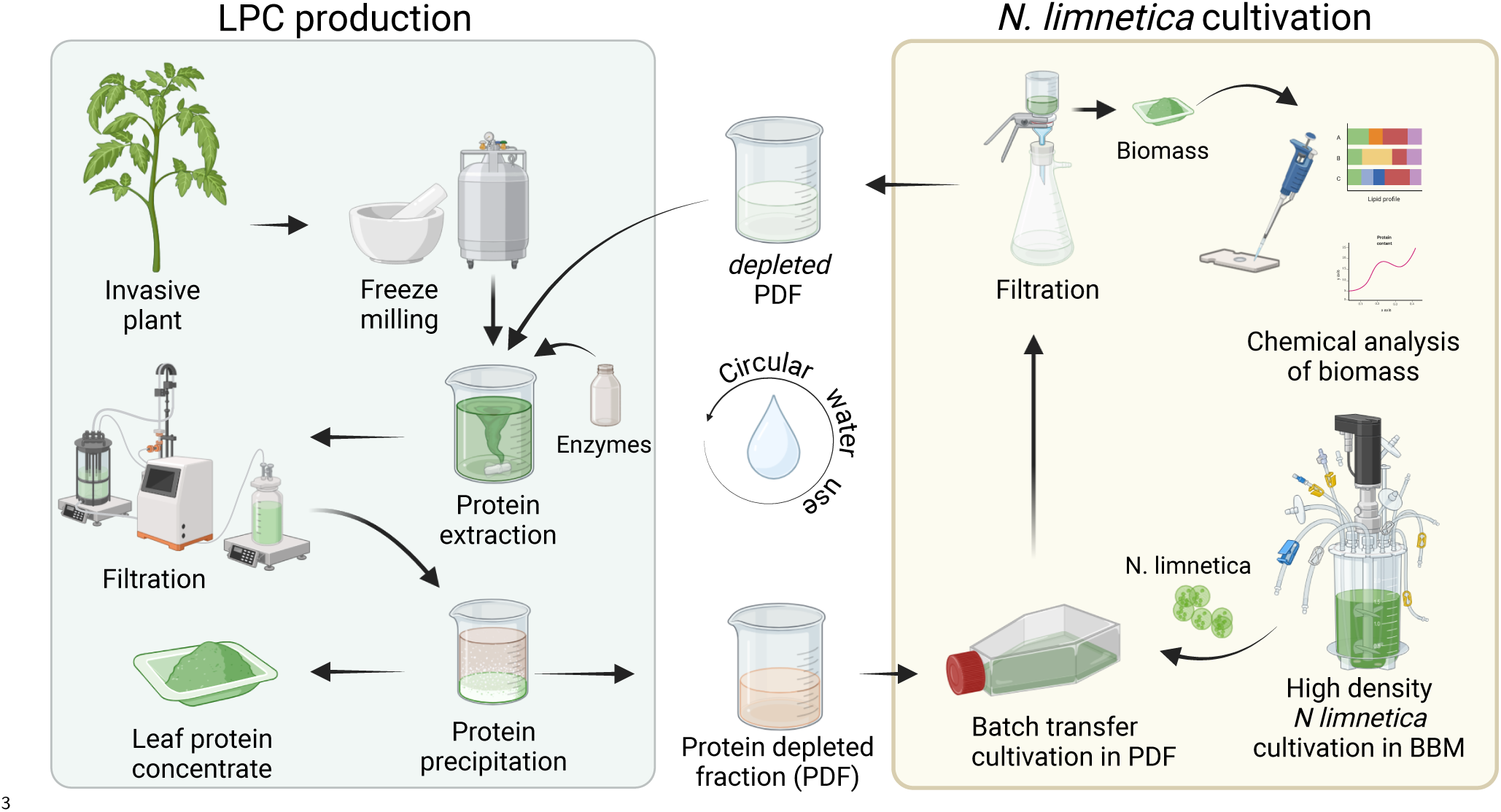

## 1 Introduction

There has been extensive reportage on sustainable wastewater treatment strategies using microalgae, owing to their robust physiologies and potential for valuable bioproducts (Catone et al., 2021; Christenson & Sims, 2011) (and the references therein). *Nannochloropsis* sp. in particular has received marked attention (X.-N. Ma et al., 2016) due to their highly lipogenic characteristics under nitrogen starved conditions (Chew et al., 2017; Levasseur et al., 2020). Based on the systematic profiling of the growth rates and lipids of all seven species of *Nannochloropsis* by Y. Ma et al. (2014), *N. oceanica* and *N. gaditana* (now *Microchloropsis gaditana*) were identified as being ideal oleaginous candidates for industrial applications. Consequently, the fresh water species *Nannochloropsis limnetica* has received comparatively poorer attention owing to its low growth rate (*µ*) of 0.07 day^−1^ (doubling time of 9 days) under standard phototrophic growth conditions, and a modest lipid production level of 74 mg L^−1^ day^−1^ (Y. Ma et al., 2014). On the other hand, the species-list noted in the comprehensive review on microalgae-based wastewater valorisation by Catone et al. (2021) almost exclusively features freshwater species, presumably to avoid the need to adjust salinity and ease down-streaming of the spent water. For example Kiani et al. (2024) claim to have successfully cultivated *N. limnetica* on whey permeate waste, while *N. oceanica* was unable to grow without adjusting the osmolarity to match seawater (∼3.2 g_salt_ L^−1^). Thus, while *N. limnetica* allows for water-water valorisation with minimal processing, its low growth rate disincentivises its widespread use.

Using *Chlorella sorokiniana* as a model organism, León-Vaz et al. (2019) could valorise acetate-rich oxidized wine lees waste in a fed-batch system. Cultivation in fed batch systems of marine Nannochloropsis species such as *N. occulata*, *N. gaditana* and *N. salina* have also been previously described by numerous authors such as Gharat et al. (2018), Martínez-Macías et al. (2018), Ocaranza et al. (2022), and Xu et al. (2004). Adjacently, mixotrophic cultivation of *Nannochloropsis* in waste streams has also been numerously published (Manzoor et al., 2021; Poddar et al., 2020). But fed batch cultivation on effluent streams has not been investigated yet for any freshwater *Nannochloropsis* species. Another consequence of the heavy focus on the lipogenic properties of *Nannochloropsis* has been the general lack of investigation on its secretome. Currently, no documentation exists on the proteomic changes relative to varying trophic conditions for *Nannochloropsis limentica*, particularly for proteases. Although changes to the intracellular protein expression levels have been reported for *Nannochloropsis oceanica* and *Nannochloropsis occulata* by Tran et al. (2016) under nitrogen paucity, differential expression in the secretome under varying trophic conditions remains unexplored and could provide the necessary context to understanding trends in protease secretion that can be applied to valorising wastewaters. However, choosing the appropriate waste stream; even for a proof-of-concept work, requires significant consideration such as cost, energy, and generating water of sufficient safety standards to warrant reuse (Acién et al., 2016; Chandra et al., 2018; Meese et al., 2022). Consequently, complete circular water reuse or repurposing in any biorefinery system is rarely demonstrated in literature, with publications generally concluding their work by measuring the nutrient depletion and only postulating hypothetical future applications, but not practically demonstrating reuse. The publication by Pinho and Mateus (2021) is a rare example where the municipal waste water treated with *Chlorella* sp. or *Tetradesmus* sp. was used to grow *Tagetes patula* L., (Marigold) in a small potted setup. Thus, to demonstrate the potential value of the proposed fed-batch style *Nannochloropsis limnetica* system, wastewater generated from the production of leaf protein concentrates (LPC) from the invasive plant Gorse (*Ulex europeaus*) will be valorised for reuse in the system.

Gorse is native to Western Europe and the Atlantic Archipelago (Rymer, 1979), has been introduced in many places around the world over the century (MacCarter & Gaynor, 1980), and subsequently became invasive resulting in the disruption of many local ecologies (Broadfield & McHenry, 2019). While numerous measures have been reported to help rid or contain the spread of Gorse, invasivory as a mode of invasive plant management has begun gaining interest (Franke, 2006; Nuñez et al., 2012), with inroads being made towards the valorisation of the invasive plant biomass through the production of leaf protein concentrates (LPC) (Iyer et al., 2022). The LPC produced from Gorse is rich in essential amino acids (EAA) (34.8 % (w/w) total protein) and branched chain amino acids (BCAA) (14.4 % (w/w) total protein) and is nutritionally complete; i.e, contains all the necessary amino acids in sufficient levels to meet all nutritional requirements without supplementation from other sources (Reynolds et al., 2022). Moreover, these amino acid levels appeared to be superior to that of soya (total EAA: appx. 21.6 % (w/w) total protein, total BCAA: appx. 7.6 % (w/w) total protein) (García et al., 1997). Additionally based on a simulated digestion model, Gorse LPC also contained residual plant phenolics that were delivered to the colon and subsequently metabolised into beneficial secondary-metabolites by the gut microbiota (Iyer et al., 2023). Thus, the LPC process appears to be an enticing way of valorising Gorse without resorting to traditional methods of removal such as herbicide use, herbivory, or incineration. However, one of the major drawbacks of the process is its heavy use of fresh water at the protein extraction stage and consequently, high volumes of effluents containing sugars, residual proteins, and plant bioactives are generated, referred to as the protein depleted fraction (PDF) (Iyer et al., 2022). The residual protein quality in the PDF had poor BCAA and EAA profiles, and consequently disincentivises further attempts to recover the residual protein using chromatography either from the perspective of nutrition or cost.

In this work, *N. limnetica* was cultivated in a fed-batch system on water from the PDF. An alternate cultivation system such as a high-density fed-batch system has been investigated using *N. limnetica* to overcome the low growth rate problem, while still generating the necessary biomass turnover to remain industrially relevant. *N. limnetica* was sought to rapidly deplete and upcycle the nutrients into valuable lipids and microalgal proteins for applications in food, feed or fuel industry.

## 2 Materials and methods

### 2.1 Gorse sample collection

Using the sampling methods described previously by Iyer et al. (2021), fresh young leaves from Gorse (*Ulex europeaus*; locally called *Aiteann gallda*; 200 g), were collected from Howth, Dublin, Ireland (GPS: 53.388, −6.065), washed, frozen at −80 °C, and freeze-dried (Lyovapor^™^ L-200). The samples were then hand milled with liquid nitrogen, mortar and pestle and stored in a cool and dark place until further use.

### 2.2 LPC production

Three batches of LPC (each in triplicate) were produced from Gorse leaf mass, which was carried out in as previously described by Iyer et al. (2022). In the first batch, the gorse leaf mass (2 g) was macerated once in water (Milli-Q^®^, USA; 40 mL) for 1 h at 40 °C and filtered to generate the first extract. In batches 2 and 3, microalgae depleted PDF (dPDF) was reused to generate the first extract (See section 2.3.2 below). The method was modified to use a higher solid to liquid ratio of 1 : 20 for each extraction step to generate sufficient volume for ultrafiltration. The plant mass was then resuspended in sodium citrate buffer (10 mmol L^−1^; pH 4.5; 40 mL) to which Viscozyme^®^ (400 µL, Novozymes, Sweden) was added and allowed to stir at 40 °C for 2 h and filtered. Volume lost to evaporation was replenished with Milli-Q^®^ water. The two extracts were pooled and subjected to ultrafiltration (2 kDa, Sartorius, USA) to achieve a retentate volume of 20 mL (4× concentrated), which was then subjected to ethanol precipitation using ethanol (7 mL, pre-cooled to 4 °C) and incubated at −20 °C overnight. The sample was centrifuged 14 000 × *g* (4 °C; 10 min; Eppendorf, Ireland) and the precipitate and supernatant were separated. The ethanolic supernatant was subjected to vacuum evaporation to recover ethanol and pooled with the remainder of the filtrate obtained from ultrafiltration to produce the protein depleted fraction (PDF, ∼60 mL). The final pH of the PDF was adjusted to pH 7.5 using sodium hydroxide solution (NaOH, 2 mol L^−1^).

### 2.3 *N. limentica* culture and cultivation

Axenic *Nannochloropsis limetica* (SAG 18.99) prepared using the method previously described by Iyer et al. (2025) was grown in standard 3N+BBM+V (40 mL) and subsequently verified using a combination of Gram’s staining microscopy, and Lyogeny Broth agar plating.

#### 2.3.1 Cultivation under different trophic conditions

*N. limnetica* was cultivated either in standard 3N+BBM+V, or Hetrotrophic growth complex (HGC). The mother culture (200 µL) was inoculated in standard BBM (40 mL) under ambient (21 °C to 25 °C) temperature conditions under constant shaking (80 rpm) with a 14 h : 10 h light cycle at 8.85 ± 0.40 W m^−2^ brightness (Barrina T5 LED Grow Lights, Full spectrum, USA). In the same conditions, mother culture was inoculated in Heterotrophic growth complex (HGC, 40 mL) for mixotrophic growth. Lastly, under the same temperature and shaking, and HGC (40 mL) wrapped in aluminium foil for heterotrophic culturing. These cultures were prepared in biological triplicate. Cultures were allowed to grow for seven days and samples (1 mL) were drawn every 24 h to measure the cell counts using flow cytometry. On day seven, the remaining culture volume was centrifuged at 3000 × *g* for 20 min at 10 °C the supernatant and cell fractions were separated, freeze-dried, and subjected to proteomic analysis.

#### 2.3.2 LPC effluent depletion using *N. limnetica*

*N. limnetica* was cultured in 3N+BBM+V (1 L) under aeration for six days (at the point of late-log phase) under the same lighting conditions noted in section 2.3.1. The culture was centrifuged at 1500 × *g* for 20 min at 10 °C. In triplicates, the microalgal biomass was resuspended in sterile filtered PDF (40 mL) generated from the LPC process (section 2.2) to reach a cell density of ∼2 × 10^4^ mL^−1^. Sample (1 mL) was drawn every 24 h and analysed for total proteins, reducing sugars, and phenolics. The spent BBM medium used to grow the *N. limnetica* was collected and refortified with the phosphate, magnesium, calcium, and trace elements stock solutions by adding 1/5^th^ of the stock volume normally required for making BBM. The medium was autoclaved, and vitamin stock solutions (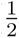 of the normal stock volume required) was added and the next batch was cultivated.

The PDF fraction from the LPC process was subjected to nutrient depletion using *N. limnetica*. After seven days of cultivation, the depleted PDF (dPDF) was filtered using a 0.22 µm filter (Millex^®^, USA) and reused in the LPC process to generate the first extract. This was done for two more rounds of LPC extraction and microalgal nutrient depletion.

#### 2.3.3 Testing phenolics removal

After the removal of *N. limnetica* cells from the dPDF in the third cycle, activated charcoal (1 mg, 3 mg, 9 mg, 27 mg, or 81 mg) was added to dPDF (10 mL) and incubated for 1 h in a roller-mixer under ambient temperature conditions (∼22 ± 1 °C). The samples were centrifuged at 10 000 × *g* for 10 min at 15 °C. The supernatant was carefully recovered as passed through a 0.22 µm filter (Millex^®^, USA) and tested for total phenolics.

### 2.4 Proximate chemical analyses

Total proteins, reducing sugars, and phenolics were measured using Ninhydrin assay, Lever assay, and Fast Blue assay respectively, as described previously by Iyer et al. (2022).

### 2.5 Lipid analysis

The microalgal biomass obtained after cultivation in the PDF was freeze-dried and subjected to lipid extraction using the method described by Bligh and Dyer (1959). Neutral lipids were profiled using gas chromatography (International Organization for Standardization, 2015).

#### 2.5.1 Flow cytometry

Cell counts were measured using flowcytometry by fixing the cells in 4 % paraformaldehyde and subsequently staining them with SYBR Green as described previously by Iyer et al. (2025).

### 2.6 *N. limnetica* extracellular protease production

*N. limnetica* was cultured for 7 days in BBM (100 mL) under standard growth conditions in triplicate. The cells were centrifuged at 4000 × *g* for 20 min at 4 °C. The supernatant was collected and filtered (0.22 µm, Millex^™^ polyethersulfone syringe filter, 33 mm diameter) and then concentrated 50× using Merck 3 kDa centrifugal concentrators (each volume concentrated at 4500 × *g*, 40 min, 4 °C).

#### 2.6.1 Detection of protease production

Protease production in *N. limnetica* was detected using two methods, both involving the use of casein agar plate. The preliminary screening step involved the use of a casein bilayer agar where the plate was prepared by first pouring casein agar (12 mL, 1.2 % w/v agar, 1 % v/w powdered skimmed milk) under sterile conditions. The plate was gently wobbled to remove gaps and spread the molten agar evenly and was then allowed to cool and set for 10 min. Then BBM medium containing 1.2 % w/v agar was similarly poured over the casein agar and allowed to cool for 10 min. Finally, 50 µL of *N. limnetica* culture was spotted at the centre, and incubated for 10 days under the standard light conditions. Milli-Q^®^ water was used as a negative control. Protease production was detected by a clear halo in the casein layer.

Extracellular protease was further confirmed by using the 50× concentrate medium after 12 days growth noted earlier, and placing 50 µL at the centre of a monolayer of casein agar (1 % w/v, 10 mL) and incubating for 16 h at 37 °C.

#### 2.6.2 Zymography for protease production

Zymogram was prepared using the protocol described by Yasumitsu (2017) with Trypsin as a positive control. This was used to assess the molecular weight of the proteases as well as test for expression during growth under varying trophic conditions (section 2.3.1) as well as in the Gorse PDF depletion cycles (section 2.3.2).

#### 2.6.3 Two-dimensional gel zymogram

Two dimensional zymogram were carried out as described by Adams and Gallagher (2004) and Wilkesman et al. (2017) respectively with minor modifications. Briefly, sample (60 µL) was added to rehydration buffer (65 µL, 8 mol L^−1^ urea, 2 mol L^−1^ thiourea, 4 % CHAPS, 1 % (v/v) Triton X, 10 mmol L^−1^ Tris Base, and 0.8 % ampholytes) and allowed to incubate in IPG strips pH 4 to 7^¶^ (Immobiline^™^ DryStrip^™^, Cytiva, Uppasala, Sweden) overnight under mineral oil in a rehydration chamber. The strips were then placed in the Ettan^™^ IPGphor^™^ II isoelectric focusing unit with the voltage program of 300 V for 30 min, 1000 V for 30 min, 5000 V for 1.5 h and 1000 V for 1 h. The strips were then placed in equilibration buffer (6 mol L^−1^ urea, 2 % (w/v) SDS, 30 % (v/v) glycerol, and 50 mmol L^−1^ Tris-HCl, pH 8.8). The buffer was gently flushed repeatedly over the strip to wash away the oil, and then drained. Fresh equilibration buffer was added and the gel was allowed to rest for 10 min. The IPG strips were then placed in standard 12 % acrylamide strength gel and sealed using agarose (1 % w/v, pH 6.8, 1 × 10^−3^ % bromophenol blue). Using a piece of a plastic comb, a groove was made in the agarose at the positive edge of the strip and 10 µL of molecular weight marker (110 kDa to 18.9 kDa, Fisher) was added. The setup was electrophoresised for 3 h at constant voltage of 109 V with a 1.5 mm acrylamide resolving gel thickness. Thiol reduction and alkylation was not performed to preserve enzyme activity.

#### 2.6.4 Predicted secreted proteases

To ensure the majority of proteases in *Nannochloropsis limnetica* reference genome were identified, all annotated proteases from Ochrophyta, the algal phylum that contains the *Nannochloropsis* genus, were gathered from the InterPro database. TargetP was used to identify the secreted proteases in the Ochorphyta protease dataset (Almagro Armenteros et al., 2019), and a homemade Python script was used to parse the results for proteases with secretion signals.

The shortlist of secreted Ochrophyta proteases and previously, identified *N. gaditana* proteases (both secreted and non-secreted) were used as queries in TBLASTN searches against the *Nannochloropsis limnetica* reference genome (GCA_001614225.1). Most of these hits did not extend to the start codon, and therefore, could not be considered for signal peptide identification by TargetP (Almagro Armenteros et al., 2019; Gertz et al., 2006). All protease sequences found in the genome are shown in Supplementary Table 5.

### 2.7 Bottom-up proteomic profiling

#### 2.7.1 Sample preparation

To check proteins in the extracellular medium, the sample was concentrated to about 10 % the original volume using a Vivaspin^®^ (3 kDa MWCO). To the concentrated sample, 4× volumes of acetone was added and incubated in −20 °C overnight. The samples were then centrifuged at 12 000 × *g* for 30 min at 2 °C (Eppendorf, Ireland). The precipitated proteins were redissolved in 200 µL resuspension denaturation buffer (pH 8.0) that comprised of urea (8 mol L^−1^, Merck, USA) and HEPES (30 mmol L^−1^, Merck, USA). Protein content was measured using BCA assay (Thermo Scientific, USA) and the concentration was adjusted to a concentration of 1.2 mg mL^−1^.

Sample was subjected to reduction and alkylation using Dithiotreol (DTT, 100 mmol L^−1^, Merck, Canada) at a final working concentration of 9 mmol_DTT_ 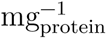 at 40 °C for 40 min, followed by iodoacetamide (IAA, 200 mmol L^−1^, Merck, USA) at a final working concentration of 18 mmol_IAA_ 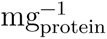 and incubation in the dark at 40 °C for 60 min.

The reduced and alkylated protein samples were dialysed against 50 mmol L^−1^ ammonium bicarbonate (50 mmol L^−1^, 50 mL) in a dialysis cartridge (Pur-A-Lyzer^™^, 3 kDa MWCO, Merck) overnight at 4 °C to remove metabolites, residual reactants and decrease the urea concentration below 1.5 mol L^−1^. The samples were then subjected to trypsin digestion for 18 h at 37 °C at a ratio of 5 µg_trypsin_ 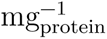 (SOLu-Trypsin, Merck)^∗^. The samples were then dried at 30 °C under vacuum using SpeedVac (SPD120, ThermoFisher Scientific), 2.5 h) and redissolved in trifluroacetic acid solution (0.05 % v/v, 50 µL). The samples were purified using C_18_ ZipTip^™^ (Merck, Cork, Ireland) according the manufacturers instructions and finally isolated in formic acid solution (0.05 % v/v, 30 µL).

#### 2.7.2 LC/MS analysis

Peptides from the sample (1 µg) was loaded onto Evotips as per manufacturer’s instructions (EvoSep). Briefly, Evotips were activated by soaking them in isopropanol, primed with 20 µL buffer B (acetonitrile, 0.1 % formic acid) by centrifugation for 1 min at 700 × *g*. Tips were soaked in isopropanol and equilibrated with 20 µL buffer A (MS grade water, 0.1 % formic acid) by centrifugation. Another 20 µL buffer A was loaded onto the tips and the samples were added on top of that. Tips were centrifuged and washed with 20 µL buffer A followed by overlaying the C_18_ material in the tips with 100 µL buffer A and a short 20 s spin.

The samples were analysed by the Mass Spectrometry Resource (MSR) on a Bruker TimsTOF Pro mass spectrometer connected to a Evosep One chromatography system. Peptides were separated on an analytical C_18_ column (Evosep, 3 µm beads, 100 µm ID, 8 cm length) using the pre-set 30 samples per day gradient on the Evosep One.

The Bruker TimsTOF Pro mass spectrometer was operated in positive ion polarity with TIMS (Trapped Ion Mobility Spectrometry) and PASEF^®^ (Parallel Accumulation Serial Fragmentation) modes enabled. The accumulation and ramp times for the TIMS were both set to 100 ms, with an ion mobility (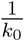) range from 0.62 V s cm^−1^ to 1.46 V s cm^−1^. Spectra were recorded in the mass range from 100 to 1700 m/z. The precursor (MS) Intensity Threshold was set to 2500 and the precursor Target Intensity set to 20 000. Each PASEF cycle consisted of one MS ramp for precursor detection followed by 10 PASEF MS/MS ramps, with a total cycle time of 1.16 s.

Data from this analysis was used to create a data-independent acquisition PASEF (dia-PASEF) method using Bruker timsControl software. The dia-PASEF settings included a mass width of 26.0 Da, no mass overlap, and mass steps per cycle of 39. The mass range was 350.0 to 1326.0 m/z. All samples were acquired using dia-PASEF.

### 2.8 Statistics and analyses

#### 2.8.1 General analyses

All statistical analyses were performed using R (4.5.0) (R Core Team, 2024). Data wrangling and visualisation was performed using tidyverse (Wickham et al., 2019), and the flow cytometry data was processed using flowCore (Ellis et al., 2024).

#### 2.8.2 Yield and growth profile

Nutrient depletion constant (k) in fed batch was calculated by modifying the standard decay formula as:

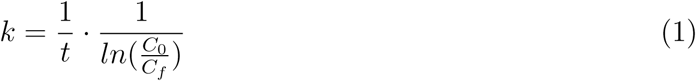

where:

C_0_ is the concentration at time point 0

C_f_ is the concentration at the final time point (day 7).

t is the total incubation time

Specific growth rate (*µ*) was calculated as:

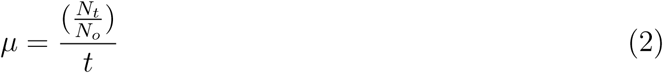

where:

N_t_ = cell number at the end of log phase,

N_o_ = cell number at the start of log phase,

t = the duration of log phase.

Yield coefficient (nutrient available per cell, Y) was calculated as:

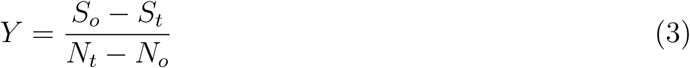

where:

S_o_ = Nutrient at the start of the log phase,

S_t_ = nutrient left at the end of the log phase.

N_t_ = number of cells at the end of log phase,

N_0_ = number of cells at the start of log phase.

#### 2.8.3 Proteomic data analysis

Bruker mass spectrometric data from the Tims-TOF was processed using Fragpipe (version 22.0) (Yu et al., 2023). Data was searched using the DIA_SpecLib_Quant which used the DDA data to build a spectral library. Quantification was carried out using DIA-NN. MS/MS spectra were matched against a Nannochloropsis database containing 33 588 entries (release 2025_03). The combined datasets for LC/MS analysis was prepared by taking FASTA databases for the entire genus of Nannochloropsis (including Microchloropsis sequences as they are yet to be distinguished in the databases) from UniProt and NCBI BLAST, concatenating them, and the using seqkit (v2.10.1) (Shen et al., 2016) to remove duplicates and correctly format the final file. The mass spectrometry proteomics data have been deposited to the ProteomeXchange Consortium via the PRIDE (Perez-Riverol et al., 2025) partner repository with the dataset identifier `PXD068602`.

## 3 Results

### 3.1 *N. limnetica* growth and nutrient depletion in Gorse PDF

The growth of *N. limnetica* on Gorse protein depleted fraction (PDF) across each water recirculation cycle along with the protein, sugar and phenolic content measurements are shown in figure 1. The duration of the lag phase appeared to shorten with each cultivation cycle. Cycle 0 had a lag phase of three days with a specific growth rate (*µ*) of 0.39 ± 0.07. Cycle 1 had a lag phase lasting two days with *µ* of 0.48 ± 0.02, and Cycle 2 showed almost no lag phase, but with *µ* of 0.35 ± 0.01. The *µ* across the growth cycles were statistically similar (F_(2,6)_ = 4.639, p=0.061) with an overall average value of 0.41 ± 0.07. Consequently, overall cell numbers on the final day of harvest were comparable at an average cell counts of 1.80 × 10^5^ mL^−1^ (∼2^16.75^mL^−1^; F_(2,6)_ = 1.479, p=0.30).

**Figure 1:**
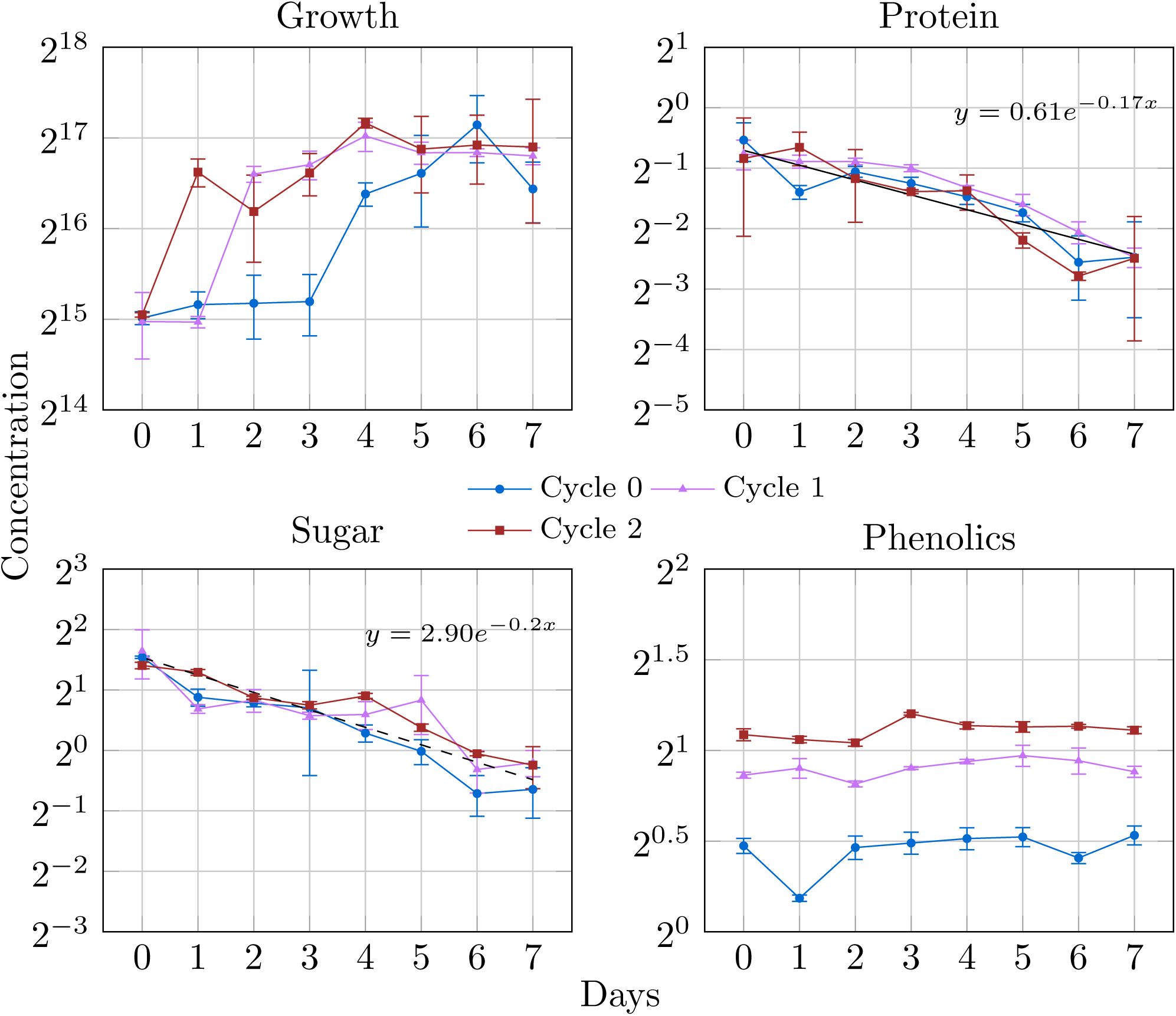
N. limnetica growth and PDF nutrient depletion profiles (n=3). The panel “Growth” shows cells mL^−1^ in the y axis and the number of days in the x axis. The panels “Protein”, “Sugar”, and “Phenolics” represent depletion or changes in the content (expressed in mg mL^−1^) over each growth cycle using Gorse PDF.

The average depletion rate k_proteins_, and k_sugar_ were 0.17 ± 0.04 day^−1^ and 0.20 ± 0.06 day^−1^ respectively. In Cycles 0, 1 and 2, the yield coefficient (Y^‡^) were 3.44 ± 1.11 × 10^−6^ mg mL^−1^ cell^−1^, 1.55 ± 0.28 × 10^−6^ mg mL^−1^ cell^−1^ and 3.62 ± 2.49 × 10^−6^ mg mL^−1^ cell^−1^ respectively for proteins. These values were statistically similar (F_(2,6)_=1.05, p=0.405) and the overall Y_protein_ was 2.87 ± 1.84 × 10^−6^ mg mL^−1^ cell^−1^. Similarly Y_sugar_ was 1.26 ± 1.10 × 10^−5^ mg mL^−1^ cell^−1^, 3.86 ± 0.77 × 10^−6^ mg mL^−1^ cell^−1^ and 7.00 ± 1.11 × 10^−6^ mg mL^−1^ cell^−1^, respectively for Cycles 0, 1 and 2. The values across the cycles were statistically similar (F_(2,6)_=0.945, p=0.44) with an overall Y for sugars being 7.81 ± 7.36 × 10^−6^ mg mL^−1^ cell^−1^. The mass balance of the proteins across the cycles is provided in Supplementary Table 1.

No depletion was observed for phenolics and consequently, it began to accumulate in the dPDF with each successive cycle. To overcome this problem at the end of Cycle 2, the dPDF was tested with increasing concentrations of activated charcoal to identify the optimal condition for phenolics removal (Supplementary Figure 1). At a depletion constant of 0.07 mg^−1^ activated charcoal, phenolics can be largely quenched from the dPDF when activated charcoal is used at a concentration of ∼32 g L^−1^ of dPDF.

### 3.2 Lipid accumulation and profile

The average biomass across Cycles 0, 1 and 2 was 1.39 ± 0.23 g L^−1^ with a total neutral lipid production of about 136.93 ± 74.86 mg L^−1^ at an average content of 92.83 ± 44.68 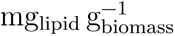. This value is about 75 % higher than the neutral lipid production provided by Y. Ma et al. (2014) under phototrophic conditions. The distribution of glycolipids, phospholipids, and neutral lipids is shown in figure 2 below and the fatty acid composition is provided in Supplementary Table 2.

**Figure 2:**
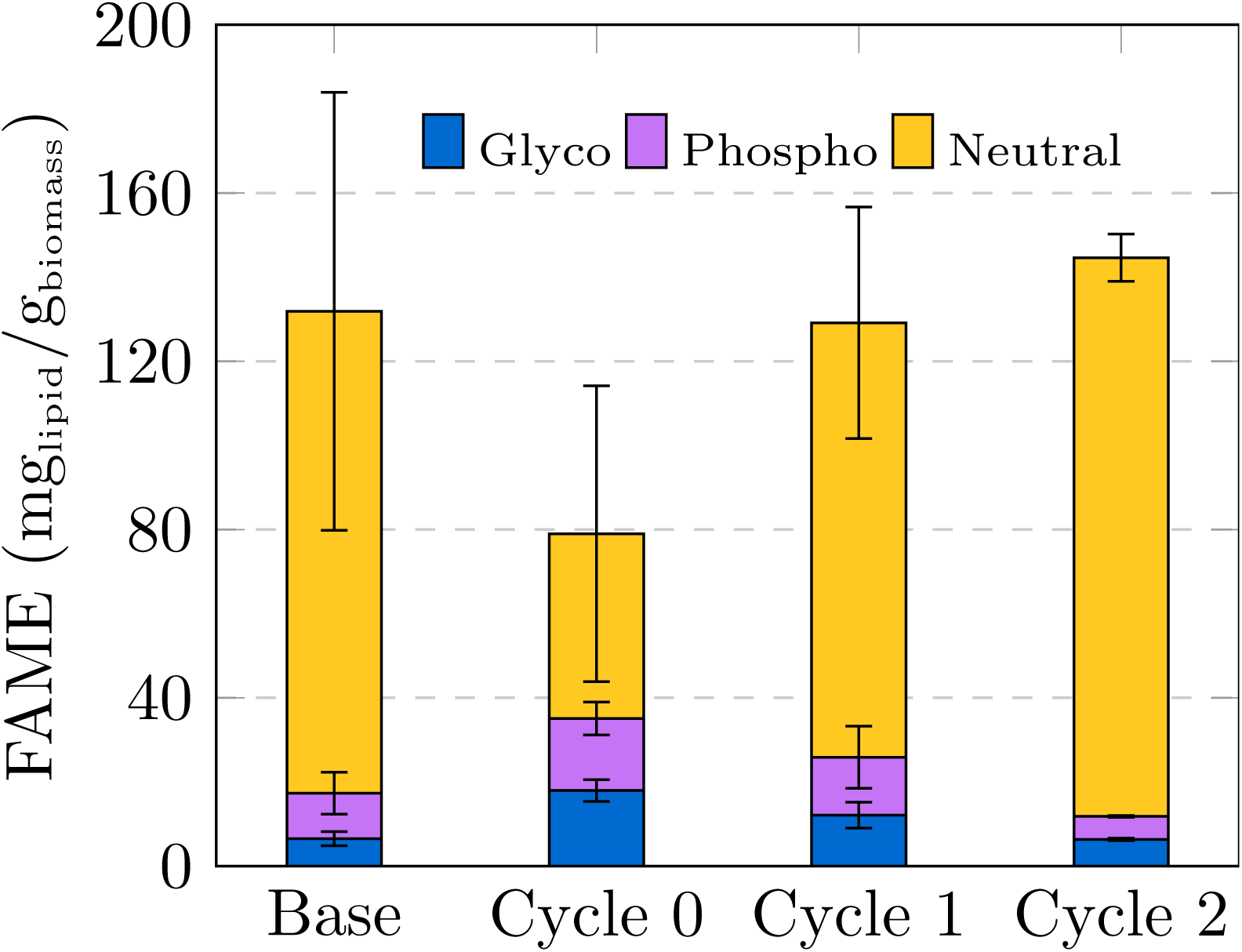
Change in lipid profiles across each cultivation cycle. Base represents the standard culture in BBM prior to the transfer of the *N. limnetica* cells to the PDF. Glyco = glycolipids, Phospho = phospholipids, Neutral = neutral lipids. Measurements were made using n=3 biological replicates.

The neutral lipid in *N. limnetica* decreased relative to the base cultivation condition under

BBM in Cycle 0. However, as the dPDF continued to be recirculated in Cycles 1 and 2, the *N. limnetica* began to show a steady recovery / increase in the neutral lipids. A one-way ANOVA comparing neutral lipid content across the batches was found to be significantly different (F_(3,8)_=7.103, p=0.012), but a TukeyHSD found that the significance was largely due to the difference between Cycle 0 and Base (p=0.05), and Cycle 0 and Cycle 2 (p=0.01). Although the neutral lipid composition appears to be the highest for Cycle 2 (132.8 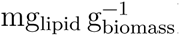) compared to the Base cultivation in BBM (114.55 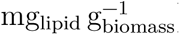), the high variation in the measurements makes comparison difficult.

### 3.3 Secretome of *Nannochloropsis limnetica*

The growth of *Nannochloropsis limnetica* in phototrophic, mixotrophic and heterotrophic growth conditions are shown in figure 3 (Panel “Growth”). Despite the high inocula density, *N. limetica* cell counts continued to double under mixotrophic conditions by three folds, while in the phototrophic conditions, the overall population doubled by about one fold. The overall cell numbers remained mostly the same in the heterotrophic conditions.

**Figure 3:**
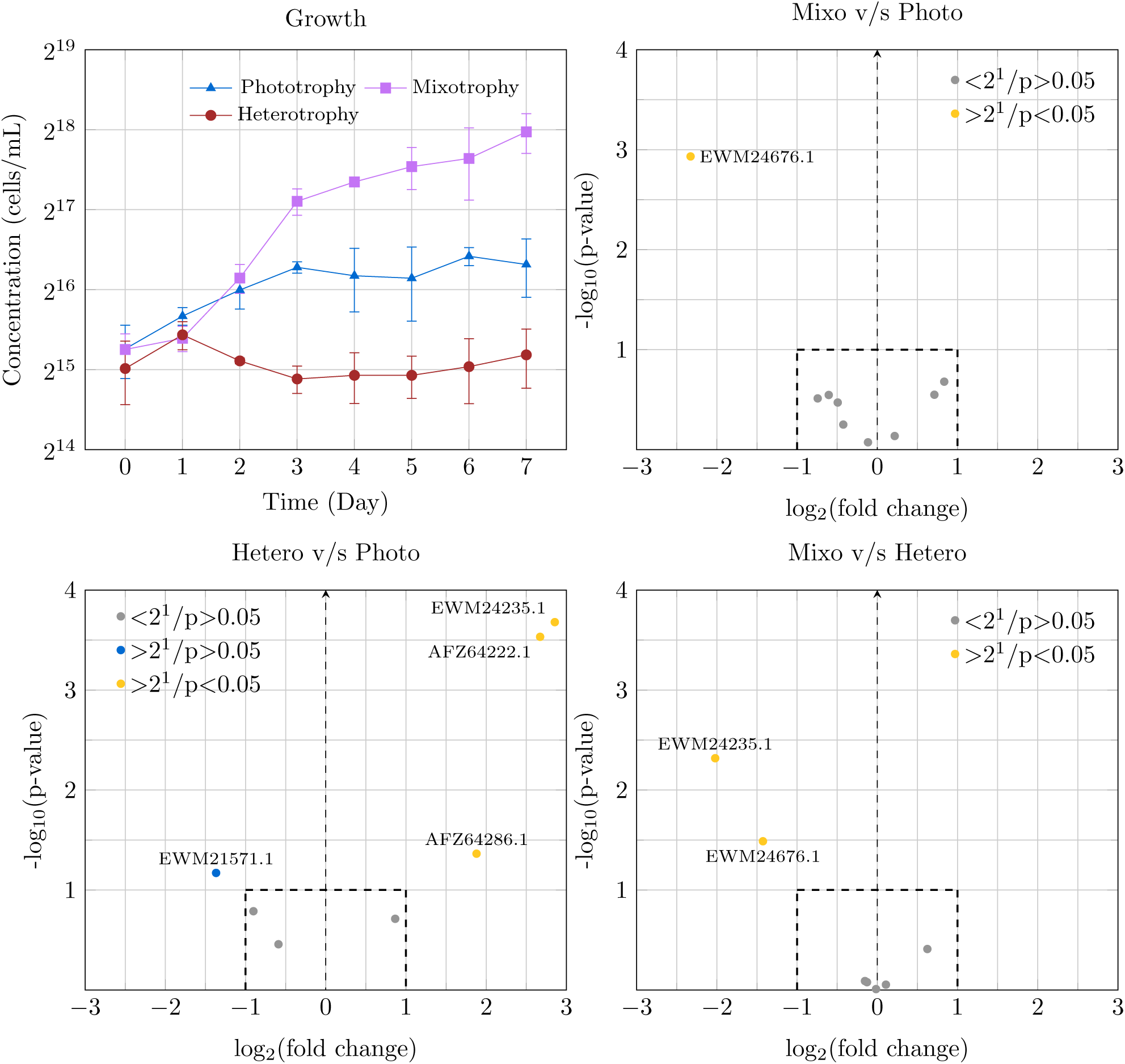
The panel “Growth” shows cell numbers over the seven-day incubation period under phototrophic, mixotrophic and heterotrophic growth conditions. Differential secretome expression under Phototrophic, Mixotrophic and Heterotrophic conditions are shown where the points marked in gray neither have an expression >2 fold, nor are statistically significant. Points marked in blue have >2 fold change, but not statistically significant. Points marked in yellow have >2 fold change and significant.

The LC/MS analysis of the secretome under phototrophic, mixotrophic and heterotrophic growth conditions are provided in Supplementary Table 5. *N. limnetica* expressed most proteins in the extracellular environment during phototrophic growth conditions rather than mixotrophic or heterotrophic conditions, with 29 out of the 45 proteins detected by the LC/MS being exclusive to phototrophy (Supplementary Figure 2). Figure 3 also shows that under heterotrophic conditions, there was a significant upregulation of `AFZ64222.1, AFZ64286.1` and `EWM24235.1` compared to phototrophic conditions, two of which are involved ATP production, while the other is crucial for nitrogen metabolism. A similar trend is observed in heterotrophy compared to mixotrophy where `EWM24235.1` and `EWM24676.1` are relatively upregulated. It appears that the histone protein `EWM24676.1` is higher in phototrophic conditions as well compared to mixotrophic conditions.

#### 3.3.1 Protease activity

Extracellular protease activity was detected firstly, with casein agar plates to test for the presence of a digestion halo either around the colony where *N. limnetica* was grown, or around the region where the concentrated supernatant was applied (Figure 5 A and B). This was characterised further with a 1D zymogram to confirm that the digestion halo was produced due to protease activity (Figure 4). The 1D zymogram was also used to provide a crude visual indication of the intensity of the enzyme activity in each experimental fraction. Lane 7 of the gel indicated lower protease activity, possibly due to lower expression under heterotrophic growth conditions.

**Figure 4:**
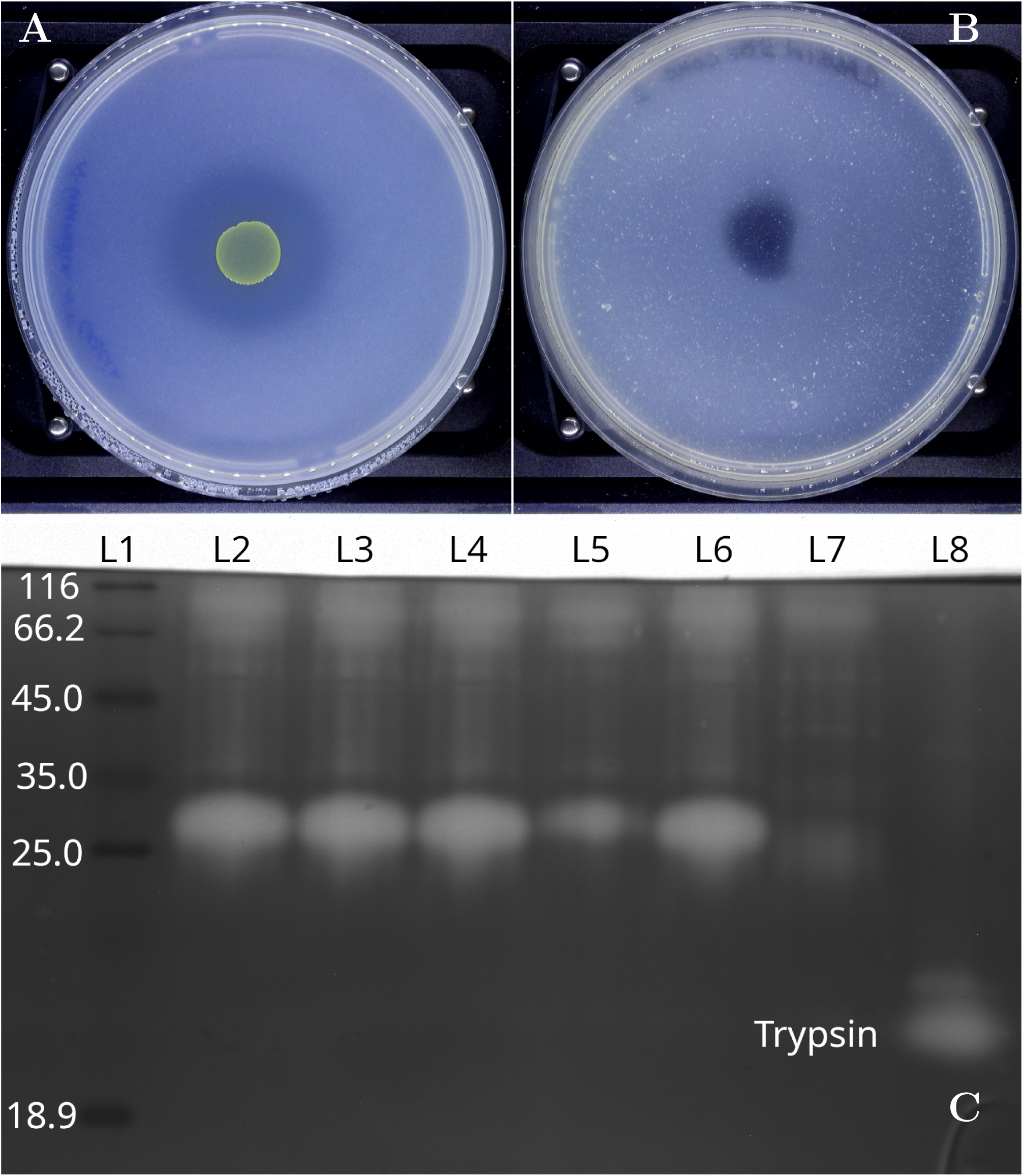
Screening and profiling of protease from *N. limnetica* supernatant (50× concentrated). Subfigure A shows the digestion halo when *N. limnetica* was grown on BBM/casein bilayer agar. Subfigure B shows the digestion halo when the 50*times* concentrated supernatant of *N. limnetica* was spotted on casein agar. Subfigure C is a casein-substrate zymogram visualising the presence of a serine protease in the 50× concentrated supernatant of *N. limnetica*. Lane 1 = Molecular weight markers, Lane 2 = Supernatant of *N. limnetica* under phototrophic growth, Lane 3 = Supernatant of N. limnetica under Mixotrophic growth, Lane 4 = *N. limentica* at Cycle 0 cultivation in Gorse PDF. Lane 5 = *N. limnetica* at Cycle 1 of cultivation in Gorse PDF, Lane 6 = *N. limnetica* at Cycle 2 of cultivation in Gorse PDF, Lane 7 = *N. limnetica* under Heterotrophic growth, Lane 8 = Trypsin (positive control).

**Figure 5:**
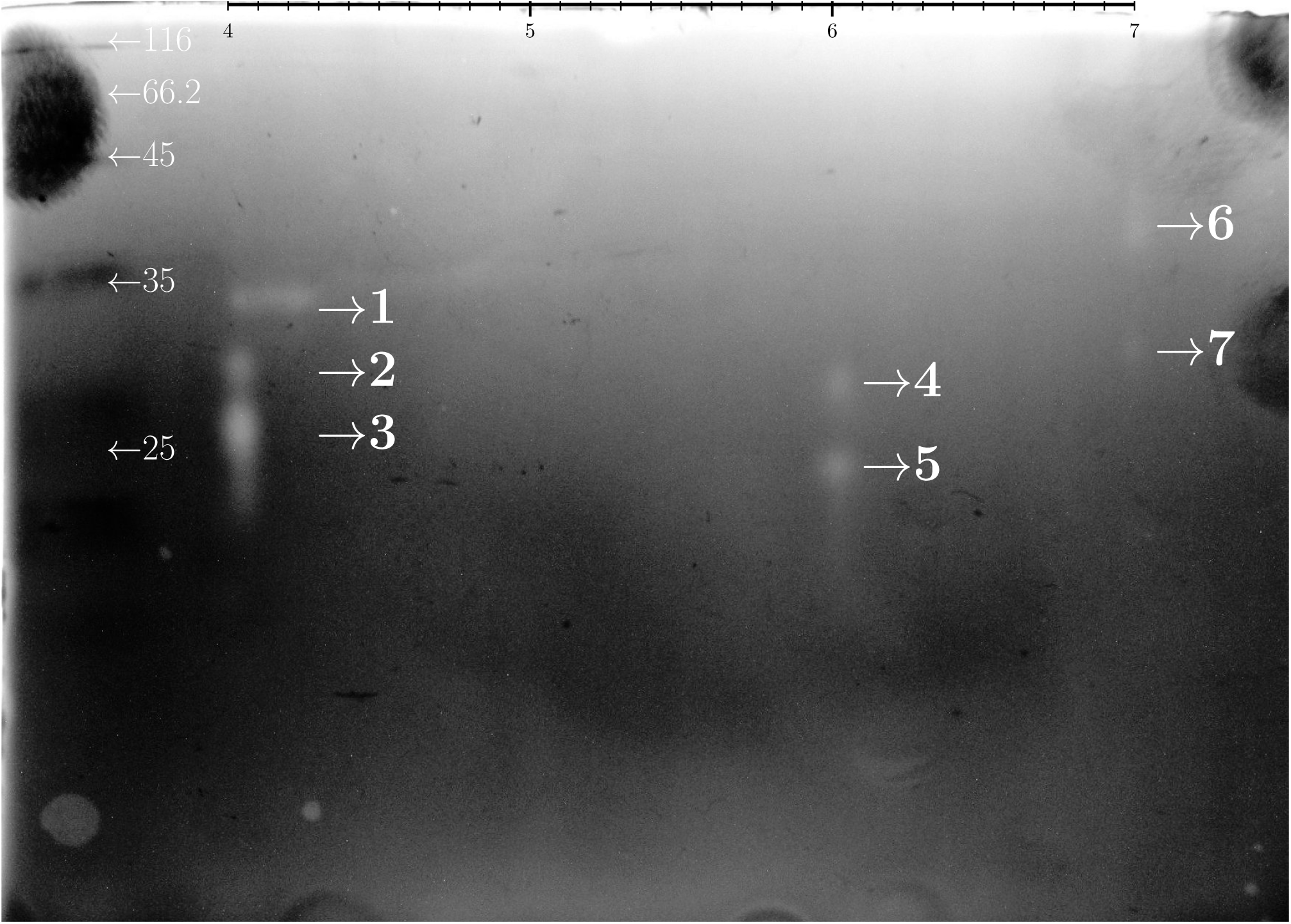
Casein substrate 2D zymogram of the 50× concentrated supernatant of *N. limnetica* showing seven negative spots indicating the presence of at least seven proteases in the sample. The scale at the top of the gel represents the pH range of the IPG strip (linear).

The 2-D zymogram (Figure 5 C) revealed seven spots labelled 1 to 7. Spot 1 (MW: 35 kDa, pI_average_: 4.3), Spot 2 (MW: 33.5 kDa, pI: 4.1), Spot 3 (MW: 29.9 kDa, pI: 4.1), Spot 4 (MW: 33.5 kDa, pI: 6.0), Spot 5 (MW: 25.8 kDa, pI: 6.0), Spot 6 (MW: 41.1 kDa, pI: 6.9), and Spot 7 (MW: 33.8 kDa, pI: 6.9). Based on the sequences from GenBank, the proteases are expected to be secreted are shown in Supplementary Table 4.

However, LC/MS analysis could not identify any of the proteases when using a database of the 31 protease sequences identified in the *N. limnetica* genome (Supplementary Table 5). Most of the protease hits occurred at the end of scaffolds and, as a result, were truncated. The only genome for *N. limnetica* is quite fragmented, and with few previous genome annotations of other *Nannochloropsis* sp., this highly limits the ability to annotate all possible proteases encoded in the genome. In addition, a lack of full-length protease sequences limited the power to identify the proteases in the secreted fractions. The protein leucyl aminopeptidase (homolog *N. gaditana* EWM21808.1 in the database) was detected in the extracellular medium under mixotrophic and heterotrophic conditions, but not phototrophic conditions.

## 4 Discussion

Previous valorisation attempts on the Gorse protein depleted fraction (PDF) to generate valuable post-biotic products such as vitamin B_12_ using fermentative methods have been mostly unsuccessful owing to the lack of carbon source to generate sufficient bacterial biomass (Iyer et al., 2024). Mixotrophic *Nannochloropsis* sp. circumvent this problem through their photosynthetic capabilities. Based on the growth profile and cell densities, it appears that *N. limentica* grows best under mixotrophic conditions, with cell numbers growing by an order of magnitude within the seven day period. Under standard autotrophic conditions, the cell numbers roughly doubled in five days; relative quicker compared to the observations by Y. Ma et al. (2014) who noted a nine-day doubling period for *N. limnetica*. The cell numbers observed for *N. limentica* under heterotrophic conditions suggest that while there was some depletion in the cell numbers, they were largely constant within the experimental period. Other indications lowered metabolism and protein secretion were observed in the zymogram (lane 7, Figure 4).

The Gorse PDF appeared to support *N. limnetica* growth quite well with an overall growth rate (*µ*) of 0.41 ± 0.07 and with the duration of the lag phase decreasing with each successive depletion cycle. While the protein and sugar content were the same after each LPC extraction cycle, the phenolics in the medium increased as it was not consumed / metabolised by the microalgae. Plant phenolics belonging to phenylpropanoid pathway such as salicylic acid, benzoic acid and ferulic acid, are known to promote growth of *Chlorella* sp., and *Nannochloropsis* sp. (Czerpak et al., 2002; Fu et al., 2021; Tam et al., 2021) and Gorse extracts is rich is these phenolic compounds (Iyer et al., 2021). Thus with each successive cultivation cycle of *N. limnetica* in dPDF, its growth was stimulated by the increasing phenolics in the medium, but limited by the depletion of the protein / nitrogen sources resulting in decreased lag-phases. This could explain the decreasing duration of lag phases in each subsequent cycle of cultivation in dPFD and help the microalgae acclimate to the medium. Thus, while the growth of *N. limnetica* was stimulated by the phenolics and sugars, the eventual depletion of nitrogen could explain the eventual limitation and plateauing of the microalgal cell numbers. Correspondingly, the lipid profile across each growth cycle was also interesting with the lipid content gradually increasing with each cycle. Phenolic intermediates of the phenylpyruvic acid pathway have been shown to increase lipid production albeit in *Chlorella* sp. and *Auxenochlorella* sp. and may have a similar effect with *Nannochloropsis limnetica* (M. Li et al., 2024; Wu et al., 2018).

Lastly, LC/MS method could identify 45 proteins in the *N. limnetica*secretome about 29 of which were exclusively expressed under phototrophic conditions. The four proteins showing significant differential expression between the pairwise comparisons of the conditions were `EWM24676.1`, `EWM24235.1`, `AFZ64222.1` and `AFZ64286.1` (Figure 3). It is interesting to note that hetetrophic growth upregulated UreG (urease accessory protein EWM24235.1) expression as it is a nickel scavenging protein necessary for the activation of urease (Myrach et al., 2017). Given that neither BBM nor HGC is supplied with nickel in the trace elements mix, the upregulation of UreG may be an attempt to scavenge ions from the medium to shift the metabolic status away from a phototrophic state to one focused on salvaging nitrogen resources with the available C : N ratio. It would also explain the absence of any of the other 29 proteins; particularly the Rubisco components that were observed under phototrophic growth as their expression will be downregulated in the absence of light. Similarly, the histone 4 protein (EWM24676.1) was upregulated in heterotrophic conditions relative to mixotrophic conditions to alter gene involved in nutrient signalling, which in tandem with the upregulation of UreG may enhance nitrogen assimilation. Even the higher presence of cytochrome c553 under heterotrophic conditions might indicate *N. limnetica* attempting to scavenge and route more ions / metals from the medium. Under mixotrophic conditions, all these proteins are downregulated compared to phototrophic and hetetrotrophic conditions as it generates the least metabolic stress (in terms of C : N ratio).

In the matter of extracellular proteases, its expression was studied to demonstrate the ability of *N. limnetica* to digest residual proteins in the medium and use it as a source of biological nitrogen. Based on figure 5 and the mass balance noted in Supplementary Table 1, it appears that the microalgae is able to secrete at least seven proteases under standard phototrophic conditions, and can actively utilise the digested proteins from the medium. The identity of the proteases however remains unconfirmed as corroborating evidence attesting to their identity could not be obtained. None of the spots in the 2D zymogram could be found in any of the samples analysed using bottom-up LC/MS. This is likely due to having insufficient coverage on the *N. limnetica*’s fragmented genome library.

Interestingly, leucyl aminopeptidase (`EWM21808.1`) was found in mixotrophic and hetetrotrophic growth conditions, but not under phototrophic conditions and wasn’t spotted in the 2D zymogram as it was carried out on extracellular proteins from phototrophic conditions. It is important to note that most of the proteins measured using LC/MS methods; especially the peptiase do not show the predicted secretion signal and thus, their expression in the extracellular environment may either be through a poorly understood mechanism, or perhaps through extracellular vesicles. The *N. limnetica* cells were grown under standard ideal phototrophic conditions inoculated at their exponential growth phase suggesting that it was unlikely this was detected owing to a large number of cells undergoing lysis / death. The cultures were axenic as well and thus, bacterial sources proteases was also technically ruled out. Such discrepancies between bottom-up and top-down proteomic results is of course well known (Z. Li et al., 2020) and comprehensive profiling of the proteases in the secretome would require further experiments as well as a more reliable and well annotated *N. limnetica* genome data.

### 4.1 Caveats and scope for future work

The work presented here demonstrates a fully circularised extraction and revalorisation bioprocess that is able to generate leaf protein concentrates (LPC) from Gorse (*Ulex europeaus*) and the high-value neutral lipids from *Nannochloropsis* sp. Previously, about 650 L of fresh water was consumed to process 32 kg of dry Gorse leaf mass to produce 1 kg of high quality LPC product (Iyer et al., 2022). The same volume can now be recirculated so the water demand can be distributed across multiple extraction batches.

Long term effects of phenolics accumulation in the dPDF on the overall process remains unclear. Within the three cycles performed herein, no interference could be observed on the protein recovery in the following LPC process and any effects noted on the microalgal growth were expected to be positive. However when nearing saturation levels of plant phenolics in the dPDF, the effects on the chemical and biochemical oxygen demand of the recirculated water may impact microalgal growth. Moreover, this process could only be demonstrated with bench-top scale volumes and higher pilot scale extraction and cultivation trials are necessary to furnish information attesting to CO_2 eq._ emissions associated to the LPC protein and the microalgal co-products.

Lastly, among the seven spots observed in the zymogram and seven proteases predicted to be secreted, only two appear to coincide at the expected pI and mol wt., and none are corroborated by through LC/MS analyses. Further investigation with size exclusion chromatography and employing synthetic peptide standards, etc. can help identify the observed proteases.

## 5 Conclusion

*Nannochloropsis limnetica* is able to help depleted nutrients from wastewater generated from the LPC production process from the invasive plant Gorse and allow for its reuse in the extraction process. It is able to accumulate higher neutral lipids in its biomass when cultivated under microtrophic conditions in waste water compared to standard medium. *N. limnetica* is able to digest the residual extracellular proteins in the effluents through the secretion of proteases although the identity of the proteases remains to be confirmed.

## Acknowledgements

Ajay Iyer is a grateful recipient of the IRC postdoctoral fellowship (GOIPD/2023/1240) and the CEA seed funding. Qinge Ma kindly helped with using the flowcytometer and Mengsong Xiao provided initial guidance and protocol with lipid measurement. Lefki Chaniotaki, Yueran Hou, Joanna Dabrinska, and provided the initial guidance and protocols for 1D and 2D gel electrophoresis, and zymogram respectively.

## Footnotes

¶ The IPG strip range was chosen based on the predicted pI values of the expected secreted proteases (Supplementary Table 4)

∗ equivalent of 50 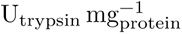, or a working ratio of 1:20

‡ the nutrient consumed per cell

