## Supplementary materials for "Protease secreting *Nannochloropsis limnetica* can valorise invasive plants through circular water-use and accumulate neutral lipids" (from another version of this preprint: version 1, 2025-11-30)

#### 1 Supplementary tables

Supplementary table 1: Mass balance of the three cycles of extraction. Values expressed in g from n=3 biological replicates.

| Fraction | Cycle | Mass | Proteins | Sugars | Phenolics |
|---|---|---|---|---|---|
| Gorse | Cycle 0 | $2.07 \pm 0.05$ | $0.15 \pm 0.01$ | - | - |
| | Cycle 1 | $2.13 \pm 0.12$ | $0.15 \pm 0.00$ | | |
| | Cycle 2 | $2.00 \pm 0.08$ | $0.15 \pm 0.01$ | | |
| Marc | Cycle 0 | $0.67 \pm 0.00$ | $0.03 \pm 0.00$ | - | - |
| | Cycle 1 | $0.64 \pm 0.07$ | $0.02 \pm 0.00$ | | |
| | Cycle 2 | $0.68 \pm 0.05$ | $0.02 \pm 0.00$ | | |
| Extract* | Cycle 0 | $1.39 \pm 0.05$ | $0.16 \pm 0.01$ | $1.13 \pm 0.05$ | $0.65 \pm 0.02$ |
| | Cycle 1 | $1.90 \pm 0.07$ | $0.17 \pm 0.02$ | $1.24 \pm 0.06$ | $0.75 \pm 0.05$ |
| | Cycle 2 | $1.54 \pm 0.03$ | $0.18 \pm 0.01$ | $1.13 \pm 0.02$ | $0.69 \pm 0.03$ |
| LPC* | Cycle 0 | $0.14 \pm 0.01$ | $0.13 \pm 0.01$ | - | - |
| | Cycle 1 | $0.14 \pm 0.02$ | $0.13 \pm 0.02$ | | |
| | Cycle 2 | $0.14 \pm 0.01$ | $0.13 \pm 0.01$ | | |
| PDF* | Cycle 0 | $1.25 \pm 0.04$ | $0.03 \pm 0.00$ | $0.10 \pm 0.00$ | $0.07 \pm 0.00$ |
| | Cycle 1 | $1.77 \pm 0.08$ | $0.04 \pm 0.00$ | $0.14 \pm 0.01$ | $0.07 \pm 0.00$ |
| | Cycle 2 | $1.40 \pm 0.02$ | $0.05 \pm 0.00$ | $0.14 \pm 0.01$ | $0.09 \pm 0.00$ |
| dPDF* | Cycle 0 | $0.41 \pm 0.02$ | $0.01 \pm 0.00$ | $0.03 \pm 0.00$ | $0.03 \pm 0.00$ |
| | Cycle 1 | $0.58 \pm 0.02$ | $0.01 \pm 0.00$ | $0.07 \pm 0.00$ | $0.07 \pm 0.00$ |
| | Cycle 2 | $0.58 \pm 0.00$ | $0.01 \pm 0.00$ | $0.08 \pm 0.00$ | $0.08 \pm 0.00$ |
| Microalgae | Cycle 0 | $0.09 \pm 0.01$ | $0.02 \pm 0.00$ | - | - |
| | Cycle 1 | $0.12 \pm 0.02$ | $0.04 \pm 0.00$ | | |
| | Cycle 2 | $0.12 \pm 0.00$ | $0.04 \pm 0.00$ | | |

Fractions marked in * include the mass from the previous cycle as the water from the dPDF fraction was used for extraction.

Supplementary table 2: Fatty acid composition of N. limnetica in each growth cycle of Gorse dPDF.

| Sample | Rep | Lipid | C16:0 | C18:2N6C | MultiC18* | C18:1N9T | C18:0 | C22:6N3 | C24:0 |
|---|---|---|---|---|---|---|---|---|---|
| | 1 | Neutral | 37.94 | 15.02 | 85.73 | 0 | 9.6 | 4.26 | 0 |
| | 1 | Glycated | 2.96 | 0 | 2.1 | 0 | 0 | 2.74 | 0 |
| | 1 | Phosphorylated | 5.44 | 2.82 | 4.61 | 3.73 | 0 | 0 | 0 |
| | 2 | Neutral | 17.68 | 4.2 | 24.38 | 6.02 | 2.95 | 0 | 0 |
| Base | 2 | Glycated | 5.45 | 0 | 0 | 0 | 0 | 1.65 | 0 |
| | 2 | Phosphorylated | 7.91 | 0 | 0 | 0 | 0 | 0 | 0 |
| | 3 | Neutral | 41.68 | 11.16 | 65.38 | 12.52 | 5.14 | 0 | 0 |
| | 3 | Glycated | 4.62 | 0 | 0 | 0 | 0 | 0 | 0 |
| | 3 | Phosphorylated | 7.98 | 0 | 0 | 0 | 0 | 0 | 0 |
| | 1 | Neutral | 8.19 | 6.03 | 9.98 | 0 | 4.13 | 0 | 0 |
| | 1 | Glycated | 8.25 | 5.58 | 6.68 | 0 | 0 | 0 | 0 |
| | 1 | Phosphorylated | 6.76 | 4.99 | 6.2 | 2.78 | 0 | 0 | 0 |
| | 2 | Neutral | 5.7 | 3.88 | 6.45 | 0 | 3.21 | 0 | 0 |
| Cycle 0 | 2 | Glycated | 7.13 | 4.94 | 6.01 | 0 | 0 | 0 | 0 |
| | 2 | Phosphorylated | 5.91 | 4.11 | 3.9 | 3.76 | 0 | 0 | 0 |
| | 3 | Neutral | 28.73 | 7.65 | 41.93 | 0 | 5.86 | 0 | 0 |
| | 3 | Glycated | 6.3 | 2.6 | 4.02 | 0 | 2.36 | 0 | 0 |
| | 3 | Phosphorylated | 4.56 | 2.56 | 2.74 | 3.1 | 0 | 0 | 0 |
| | 1 | Neutral | 42.22 | 11.66 | 72.11 | 0 | 7.5 | 0 | 0 |
| | 1 | Glycated | 3.64 | 0 | 2.89 | 0 | 2.29 | 0 | 0 |
| | 1 | Phosphorylated | 3.1 | 0 | 2.41 | 0 | 0 | 0 | 0 |
| | 2 | Neutral | 79.9 | 4.55 | 5.13 | 2.79 | 4.37 | 0 | 0 |
| Cycle 1 | 2 | Glycated | 6.69 | 4.45 | 3.79 | 0 | 0 | 0 | 0 |
| | 2 | Phosphorylated | 6.02 | 3.54 | 2.59 | 7.61 | 0 | 0 | 0 |
| | 3 | Neutral | 66.08 | 3.43 | 3.88 | 2.3 | 3.94 | 0 | 0 |
| | 3 | Glycated | 5.43 | 3.79 | 3.29 | 0 | 0 | 0 | 0 |
| | 3 | Phosphorylated | 4.88 | 3.03 | 2.2 | 5.95 | 0 | 0 | 0 |
| | 1 | Neutral | 42.3 | 11.41 | 70.86 | 0 | 7.82 | 0 | 2.42 |
| | 1 | Glycated | 3.8 | 0 | 2.23 | 0 | 0 | 0 | 0 |
| | 1 | Phosphorylated | 3.09 | 0 | 2.6 | 0 | 0 | 0 | 0 |
| | 2 | Neutral | 43.14 | 11.68 | 72.31 | 0 | 7.88 | 0 | 2.15 |
| Cycle 2 | 2 | Glycated | 3.94 | 0 | 2.45 | 0 | 0 | 0 | 0 |
| | 2 | Phosphorylated | 3 | 0 | 2.36 | 0 | 0 | 0 | 0 |
| | 3 | Neutral | 40.18 | 11.09 | 68.39 | 0 | 6.8 | 0 | 0 |
| | 3 | Glycated | 3.66 | 0 | 2.95 | 0 | 0 | 0 | 0 |
| | 3 | Phosphorylated | 3.04 | 0 | 2.31 | 0 | 0 | 0 | 0 |

* MultiC18 = C18:1N9C/C18:2N6T/C18:3N3

Supplementary table 3: Dataset of the proteins identified in the extracellular environment of N.limnetica under phototrophic, mixotrophic and heterotrophic conditions. Biological replicates (n=3).

| |
|---|
| Data available in `Peptides_full.csv` |

Supplementary table 4: Protein Accession Data

| Accession | Amino acid sequence[3] | Length[2] | MW (Da)[1] | pI[1] |
|---|---|---|---|---|
| AFGP01000834.1 | MRSSQILTLAVALCGSFSMTLTSA–YQSPSFAPAPSRASRWMLGSRSAGVAPTATFNPPAASARGARGGAGSLVMMPIGVPKVAYRVPGSGSADWVDIYNRLYRERIIFLGQEIDDDYANQIIGVLLYLDQEDSSRPIYMYINCPGGSVIAGLAIFDVMQLIRSEIVTINIGMAASMASFLLAAGKKGKRLALPHSRVMIHQPMGGAQGQAEDIKVEAAQILRIKKNLVGMYSKMTGRTTEQIVIDLDRDNYLSAQEALDHGLIDEIVELEKAMEREENPNVNLDPVRAFAGVAKDDDPLKYLP | 348 | 30353.87 | 5.72 |
| AFGL01004582.1 | MRFLSLLLLGFVATATVAT–TTTISTSTFDNEDPSSSPLLPLFRAWMEHHNISIPKAHFAHRLRVFAANHAFILAHNAQLHPRPSYFLGHNRFSHLTFDEFRTMHLGWTRPTEALPPSFPPSFENEDERTTTTTRALRGSIAPADDNNTNSVVGERYPSYTFPLPAWLPHPDPSSEDPSSLPASIDWVSKGAVTPVKDQGSCGSCYSFSAVGAVEGAYFLANGKLLELSMQEIVDCDDLDGGCQGGEMQQTFKWMKSNGGLCLLSDYPYVSGTTAEEQKVCSTCPLVPGTAPESWTTVRRNSMEGLKEAVAKGPVSVAVGASRAWMFYESGVFDGACEEELNHGVLCVGYGGGEGEEAGEEG | 361 | 37133.24 | 4.91 |
| AFGL01008529.1 | MRSSQIVALAVALCGCLVSA–YRSPSFAPAPSRASRWMLGSRSAGVAPTATFAPPAASARGARGGAGSLVMMPIGVPKVAYRVPGSGSADWVDIYNRLYRERIIFLGQEIDDDYANQIIGVLLYLDQEDSSRPIYMYINCPGGSVIAGLAIYDVMQLIRSEIVTINIGMAASMASFLLAAGKKGKRLALPHSRVMIHQPMGGAQGQAEDIKVEAAQILRIKRNLVGMYSKMTGRTTEQIVIDLDRDNYLSAEEALAHGLIDEIVELEKAMDREENPNVSLDPVRAFAGVGKDDDPLKYLP | 345 | 30284.82 | 6.01 |
| AFGL01011628.1 | MKWVFQPSFVLPCAFFLAVVLTFLRS–APQKVEWNPHTYCPPLSTFLQIDVLSRLLKDRILLLGQQVDDEVANILVAQLLYLAQEDKEKDITLYINSPGGSVSAGLAIYDAMQFVPCDVATVCFGMAASMGAFLLGAGAKGKRRSLPNARIMIHQPLGGAQGQAADIEIQVGVQAKESLFVRFFLNSYISEYNGQPIDKIQEDCDNAFSTHSHLSIPPSPPPSFYHRLRRSCLSAPCSTATSASTPASRSTRSRRIATVIFS | 313 | 25571.26 | 7.81 |

Continued on next page

Table 4 continued from previous page

| Accession | Amino acid sequence | Length | MW (Da)* | pI* |
|---|---|---|---|---|
| AFGL01011628.1 | MRAVILLVAALVLATLSEAQA–<br>FIAPAARFNAASSRVASSKKTVMMSSPDLWYPGMKWD<br>GRGLRMSPDQEPEVDPRRKEEQKPERFPVRLPVIETQ<br>TGQIDVLSRLLKDRILLLGQQVDDEVANILVAQLLYL<br>AQEDKEKDITLYINSPGGSVSAGLAIYDAMQFVPCDV<br>ATVCFGMAASMGAFLLGAGAKGKRRSLPNARIMIHQP<br>LGGAQGQAADIEIQQAKEILFVRSLLNSYISEYTGQP<br>IDKIQEDCDRDFFMTPEEAVAYGMIDSIVPTKTSHLK<br>KPTMPALLW | 332 | 29618.17 | 5.57 |
| AFGL01021278.1 | MAHRLSFLLLPLLLPLLTSA–<br>FHAPMPTRRTSSSWVVSGPRSSSSSSSLSGLSSGNGRRPPLP<br>PSYLAFPHLFASGNNGEGEEEEQERKEEDGSLVDKAKSFFA<br>DPENRKDTTLFAVSFIVSLAIRTFIVEPRYIPSLSMYPTFL<br>VGDQLAVEKVTKTYKNYERGDVVVFNPTQGYQEYVSQDP<br>YITDKSKINEALIKRIIAKEGDVVEVKDGLLFINGAAQD<br>EKYIAEGPAYVWGPNRVPDGMYMVLGDNRNHSLDSHIW<br>GFLPKENIIGKAICKYWPPWRLGLVEGPFQQAS | 350 | 30446.3 | 5.7 |
| AFGL01021448.1 | MARPTTPLFVLALLALLLSHCPTQA–<br>ESWFPPERCRGGTLWQDFVARHPYAAAPSSHAHTWDAP<br>IDHFNADDTSTYKQRYFLDAQYFQPSANGTGLVFLMI<br>GGEGTLQGPPGGFVAELAKEHNALLVSLEHRFYGES<br>FPNHDMSTAALSQYLTVDQALADLAAFSDYLQSEKFP<br>DTPLRFFAIGGSYPGALASWYRIAYPDKTLGSLSSSG<br>VVNAILDFPDFDRYVALAIGEPCAGRLRKVTRAFEDAL<br>RVPAAACVTKGLFGCDCDMWDADFFYMLADAAAMVDQ<br>YGRKEELCSFLERLEEGEEGREEEGAPSDFELVWGFA<br>NFTNSYYGPQFASSCFYDSRCAGDVSKATLNERSWRW<br>QKCSELAYFQVAPQEEDEEGREEGSLRSGEVTLEYHL<br>EQCRRIFGKGVGEPKVDEINARFGGARPKATRVFYSD<br>FSDDPWQAASVLEEEEGEGG | 478 | 47642.66 | 4.56 |
| AFGL01023503.1 | MTTSSTTTTPATTRRWRPCLLPSVASLLLLLVTLAPTG[illegible]A–<br>FFFGLPTSSSSTCHAAAAAAAAAPTPSSSIATSSSSFP<br>WRACSATSTARPTSTTRTYATAAAGGNDEEEDGLPDES<br>DAPEPSATTDTWRSNGQEPPPPPAKHPTQLDLTGLPAK<br>VMGSVFKLKVTSCPPLFRRPWSRQPTSTSFSTAWAYDV<br>KEKLLLTNSHCVMDAVVINVKKPGNSNLFKAEVLGKSD<br>HSDLALLRVPDDGFWEGVEAIRMEPEDVKLGELVTVIG<br>YPRGGEKICLTKGIVSRLHFNGEYLSIQIDAVRVPGRG<br>ERERGRE | 389 | 29246.81 | 6.01 |

[1] MW and pI are predicted values based on sequences without the signal peptides.
[2] Length is the complete protein peptide length including the signal peptide.
[3] The – in the sequence demarcates the end of the signal peptide and start of the expressed final protein.

Supplementary table 5: Proteases in N.limnetica

| Accession | Status | Contig end | Closest Homolog | Protease Sequence |
|---|---|---|---|---|
| AFGL01000090.1 | Partial | Y | XP_005854463.1 | PPSLPPSTPQTVTKAQL-TIAED GRKVGYLK-LKEFNSLAKEKVRD ALSSLEE-QGAEAYIIDLRGNPG GSVQAAVSVASLFLPED-KIVTFI QDATGNRLPLK-TQGAAALPPS DPIVVWT-DRKSASASEILASAL HD-NCRAVLMGQRTFGKGLVQ GVFGLTDGKGLVMT-VAKYETPR GASIQGT-GILPDIEAVIPSNILGD QD-VSVVRKEDFGVKDGMCLEAL ASK |
| AFGL01001552.1 | Partial | Y | XP_005854463.1 | MPDSIICRRGEASRLRCAATR SCLPFDYFSKDAPHRTHTPLH RQQASQSCLLSTRRMAT-SAT RRRQSGSAMTST-SPLSLLASL LLSLALLLVL-RPPQAVDAFRIRS SALSAP-STSCPRAGGWTRRGT AS-RTMTMRQQGPNQGDDVKK QQQPPSPAHLWVLETLHRTSK LAPGFLGLVLGARPLFAVAP-PAY AAPEAATTTTTAAP-PASSVTAP TFSSDQAASDS-DDLFDEVWKLT SKFYL-DRSFGGSDWGKARMDLR AQGKLPGDSASETKATKKL-LKKL GDKYTQLL-SPYMYTEMSRFDPI GAGFML-SIDDDGYFCVSSDPREG SVAAAQKVQKGDKILEIEG-VAIKGK SVFDAVDLITKED-KADVRLTIQSK LDPGAAPRL-LVLPRDFNAVTNPG EGRKG-GREGGKGGGGM |

Continued on next page

Table 5 continued from previous page

| Accession | Status | Contig end | Closest Homolog | Protease Sequence |
|---|---|---|---|---|
| AFGL01002145.1 | Partial | Y | XP_005856251.1 | MRAPARVVFGEAVKLITEA-<br>CLHAV QGYNRSDRHGF-<br>FVASAEGTPAAG TAG-<br>VAAGGEGGYCLLNLH-<br>NDDVHT YNQVTDALL-<br>RLRFNAASAQTLTER<br>VDTQGHAVVKRGKATELKPMLDSL<br>ASHGLLVSLVDEYQINRGER-<br>AALLL GWLTGLSTQVDGLAR-<br>IVAELLAEAC PLGLSQL-<br>GYREGTRWGTPLVRIYQP<br>APDDASLVLSTEDAKLNLSEG-<br>RLSSR LALLMMCDFFLLK-<br>TLRTELHALYLH LLV-<br>DAHFKPALAKALTWCY-<br>PALNSLY SRGVGTHDDSLFS-<br>FSVQLYTTPSIVR WLATT-<br>TADEEELGSKGLLRVLTEAVC<br>QAFKIAGWREGQLQPLPES-<br>FWEHPLL QHRRYAHLL-<br>RDIDYALQTEGTAAAVL<br>RQAYPGPGPHPLPLWVSVL-<br>GALQHAD GTMRKTGAHLE-<br>FESTVWTQVSKQEGG REG |
| AFGL0100282.1 | Partial | Y | XP_008892743.1 | MATLRSLFPVLHTTI-<br>ASYIDANDIFNV RSAAR-<br>WCQDVYGPQLMTKQF-<br>SIRIQP QAKKAKRAT-<br>PAAQAKKARAGTTTAS<br>TATAPKTTMRPPETT-<br>TFEPEVEATGAE ARVEGPP-<br>GAYSARHDPMWEALAAIE<br>ETLPQRSEARLRRDL-<br>GAMIAVGGKGM LIRM-<br>RMAPLTEEGQEEIHRLL-<br>WQGR GTDVVVSGFNVN-<br>MRRQDIRTCAPYQ WL-<br>NDEVINFFMKLLKERD-<br>DALVAAAV ARGEERRKN-<br>WFFNSFFITKFLDDDKG<br>YKYANVKRWSRTAGDLF-<br>SYDKVFFP HIIGNSHW-<br>CLVVIHMQDKKIEYLDS<br>LVGTKRDYLPHLLQYLQDE-<br>HQDKKK AALD |

Continued on next page

Table 5 continued from previous page

| Accession | Status | Contig end | Closest Homolog | Protease Sequence |
|---|---|---|---|---|
| AFGL01003586.1 | Partial | Y | XP_005855160.1 | GEEGGEEGEEYWKVKN-<br>SWGTGWG MEGYILLKRDG-<br>GDKKVACGILQDASF PILP-<br>KPSSSSSSSSSTTSSTSNNDN<br>NIISSSTTPSSPLQGSA-<br>TARDCGGGTA AFSSI-<br>TITPSSIEKNQLVTVLAS-<br>GVLTK PTLSGSYSLSV-<br>TYEGAQLYSHAGDAC<br>GNETVSLPLNAGTVHVFG-<br>FACPANPG PVAYGLDVTLPSI-<br>APSGGYLISIKGKDQ DGAALV-<br>CIEAKLEL |
| AFGL01003881.1 | Partial | Y | XP_002184343.1 | QSVRYTPPSQGGEG-<br>GKEDEATKD SKQKR-<br>RAASFLKEGGGGGRGGFE<br>ANDVPESERQIFYTKG-<br>GRVVRDGG GVEADIK-<br>VAAPQASVLEVS-<br>LAQQG VFFDFANEWAK-<br>SHTYSLGEGLKEG<br>GPALVSEETY-<br>DDFLQFVKKKEKEG<br>SLRLDLLFSGSLEPLK-<br>NALKATEYS KAGKELEGL-<br>REAVREEVQKEFLTEKK<br>QIKEEVEQAILARYLPESMLR-<br>RLAL EGDPQVKAAIEAVREG-<br>GREGG |
| AFGL01006177.1 | Partial | N | XP_005649773.1 | RRFFSTLSLLPVAITVNEMVG-<br>GLMF VDGRSMQPTLNS-<br>DNNGGLNDSRDLVL VAKW-<br>PIKVWRQYRKGDVVVLR-<br>SPIDP ETKVIQRLT-<br>GTEREWRRRRTGVRQEP<br>VPEGFCWVSGDNPLFSED-<br>SQTTYG PIPLALIEGRVSY-<br>ILWPPSRIGPVPKQ IPVN-<br>GRQPPGRVPPQEV |

Continued on next page

Table 5 continued from previous page

| Accession | Status | Contig end | Closest Homolog | Protease Sequence |
|---|---|---|---|---|
| AFGL01008011.1 | Partial | Y | XP_008892743.1 | RGRDVVVSGYGVYMQRM-<br>SIRRCGLC KWLNDEV-<br>INFFMQLLKQRDDALVAAA<br>VARGEERLKNWFFSSY-<br>IMSKLLDDDK GYKYAN-<br>VKRWSRAAGDLFTYD-<br>KVFF PVNIGNSHWCLAV-<br>IHMQAKKIEYLDS LIGT-<br>GREYLPHLLQYLQDE-<br>HQDKKQ AALDMSG-<br>WELVYSDPQVVPQQANGS<br>DCGCFVSAFAYLLSEGLPIKY-<br>ISPADMP AFRQRLVM-<br>SIMKKSLTW |
| AFGL01008078.1 | Partial | Y | XP_005854401.1 | RGGEGGGAQLVRLTTL-<br>PANGILHS DQHDLMPID-<br>NDRGTRPLRRFAA LLLL-<br>RPLPDTVNVDLSTPPAVSL<br>QWDEEEEGEEEGLPLREL-<br>SRD KMFWERVFALVAPEP-<br>SLVGDA WALLQMV-<br>PAIPLIEESIRSLGGA LSLP-<br>NSSSTSSSSSTSTGGVDGG<br>ASNTSSGSSTTTSSSSSSSNE<br>IRTWEALLHGRSPMRFLYNLQ<br>LIQDKILFPSTDPYSSPSS-<br>RASS LRWELAFVKMG-<br>GLAYLVEVLAH WDIAS-<br>ESKNPTLDKNTLSALIQC<br>IGNFMPPKQQEEEEGRVF-<br>GRGGGGG |

Continued on next page

Table 5 continued from previous page

| Accession | Status | Contig end | Closest Homolog | Protease Sequence |
|---|---|---|---|---|
| AFGL01008086.1 | Partial | Y | XP_005855948.1 | SLPPSLPPSSRYIIPVEVVHHF<br>LEDFRRHGQDMGA-<br>CLQGFELQ DLTNTALRES-<br>VAGNHTGGLVID VAGWT-<br>NAAEHLKAGDVVLSIDG<br>QKLQNDKTIPFKDSDYINFE-<br>FLM SYHFVGDMIKMEVL-<br>RRDQPLEDEE GAALP-<br>PLATFHTRNLTWEMQKRHP<br>ILPSRPQNQGFLMVG-<br>GLLFLPL SWEILDP-<br>DRSTRALRSFCECVRR<br>VYADEEVVVLSEIF-<br>PHAINDGYLP PRLTRVLSFN-<br>DEPIRNLTHLTKMI EDC-<br>QGDWMRFKVDSNLQQ-<br>TIVLD ARENAVWNATRE-<br>ICQVYSVPYYK PLPPM-<br>RKEGEEGAEEEQEVPPWM<br>RRRGGGGGGMGWGLEGGE-<br>GRW |
| AFGL01004582.1 | Partial | Y | XP_005855160.1 | MRFLSLLLLGFVATATVATTT<br>TISTSTFDNEDPSSSPLLPLF<br>RAWMEHHNISIPKAHFAHR<br>LRVFAANHAFILAHNAQLHP<br>RPSYFLGHNRFSHLTFDEFR<br>TMHLGWTRPTEALPPSFPP<br>SFENEDERTTTTTRALRGSI<br>APADDNNTNSVVGERYPSY<br>TFPLPAWLPHPDPSSEDPS<br>SLPASIDWVSKGAVTPVKD<br>QGSCGSCYSFSAVGAVEGA<br>YFLANGKLLELSMQEIVDCD<br>DLDGGCQGGEMQQT-<br>FKWMK SNGGLCLLS-<br>DYPYVSGTTAEE QKVC-<br>STCPLVPGTAPESWTTVR<br>RNSMEGLKEAVAKG-<br>PVSVAVGAS RAWM-<br>FYESGVFDGACEEELNH<br>GVLCVGYGGGEGEEAGEEG |

Continued on next page

Table 5 continued from previous page

| Accession | Status | Contig end | Closest Homolog | Protease Sequence |
|---|---|---|---|---|
| AFGL01008529.1 | Partial | Y | XP_002292583.1 | MRSSQIVALAVALCG-<br>CLVSAYRSP SFAPAPSRAS-<br>RWMLGSRSAGVA PTAT-<br>FAPPAASARGARGGAGSLV<br>MMPIGVPKVAYRVPGSGSAD-<br>WVD IYNRLYRERIIFLGQEID-<br>DDYANQI IGVLLYLDQEDSSR-<br>PIYMYINCPG GSVIAGLAIY-<br>DVMQLIRSEIVTINI GMAAS-<br>MASFLLAAGKKGKR-<br>LALP HSRVMIHQPMG-<br>GAQGQAEDIKVE AAQIL-<br>RIKRNLVGMYSKMT-<br>GRTTE QIVIDLDRDNYL-<br>SAEEALAHGLID EIVELEKAM-<br>DREENPNVSLDPVRA<br>FAGVGKDDDPLKYLP |
| AFGL01011628.1 | Partial | Y | XP_002184857.1 | MKWVFQPSFVLPCAF-<br>FLAVVLTFL RSAPQKVEWN-<br>PHTYCPPLSTFLQ IDVL-<br>SRLLKDRILLLGQQVD-<br>DEVA NILVAQLLY-<br>LAQEDKEKDITLYINS PG-<br>GSVSAGLAIYDAMQFVPCDVA<br>TVCFGMAASMGAFLLGA-<br>GAKGKR RSLPNARIM-<br>IHQPLGGAQGQAAD<br>IEIQVGVQAKESLFVRF-<br>FLNSYISEY NGQPIDKIQED-<br>CDNAFSTHSHLSIP PSPPPS-<br>FYHRLRRSCLSAPCSTATS<br>ASTPASRSTRSRRIATVIFS |

Continued on next page

Table 5 continued from previous page

| Accession | Status | Contig end | Closest Homolog | Protease Sequence |
|---|---|---|---|---|
| AFGL01021278.1 | Full length | N | XP_005855049.1 | MAHRLSFLLLPLLLPLLT- SAFHAP MPTRRTSSSWVVS- GPRSSSSSSS LSGLSSGN- GRRPPLPPSYLAFPHL FASGNNGEGEEEEQER- KEEDGSL VDKAKSFFAD- PENRKDTTLFAVSF IVS- LAIRTFIVEPRYIPSLSMYPTFL VGDQLAVEKVTKTYKNY- ERGDVVV FNPTQ- GYQEYVSQDPYIT- DKSKIN EALIKRI- IAKEGDVVEVKDGLLFIN GAAQDEKYIAEGPAYVWGPN- RVP DGMYMVLGDNRNHSLD- SHIWGF LPKENIIGKAICKY- WPPWRLGLVE GPFQQAS |
| AFGL01021448.1 | Partial | Y | XP_008907881.1 | MARPTTPLFVLALLAL- LLSHCPTQ AESWFP- PERCRGGTLWQDFVARH PYAAAPSSHAHTWDAPID- HFNAD DT- STYKQRYFLDAQYFQPSANGT GLVFLMIGGEGTLQGPPG- GFVAEL AKEHNAL- LVSLEHRFYGESFPNHD MSTAALSQYLTVDQALAD- LAAFS DYLQSEKFPDTPLRF- FAIGGSYPGA LASWYR- IAYPDKTLGSLSSSGVVN AILDFPDFDRYVALAIGEPCA- GRLR KVTRAFEDALRV- PAAACVTKGLFG CDCDMW- DADFFYMLADAAAMVD QYGRKEELCSFLERLEEGEE- GREEE GAPSDFELVWG- FANFTNSYYGPQFA SS- CFYDSRCAGDVSKATL- NERSWRW QKCSELAYFQ- VAPQEEDEEGREEGSL RSGEVTLEYHLEQCR- RIFGKGVGEPKV DEINARFG- GARPKATRVFYSDFSDDPW QAASVLEEEEGEGG |

Continued on next page

Table 5 continued from previous page

| Accession | Status | Contig end | Closest Homolog | Protease Sequence |
|---|---|---|---|---|
| AFGL01023503.1 | Partial | Y | XP_005704064.1 | MTTSSTTTTPATTRRWRP- CLLPSVAS LLLLLVTLAPT- GRAFFFGLPTSSSSTCH AAAAAAAAAPTPSSSI- ATSSSSFPWRA CSAT- STARPTSTTRTYATAAAG- GNDEEE DGLPDESDAPEP- SATTDTWRSNGQEPP PPPAKHPTQLDLT- GLPAKVMGSVFKLKV TSCPPLFRRPWSRQPT- STSFSTAWAYDV KEK- LLLTNSHCVMDAVVIN- VKKPGNSNL FKAEVLGKS- DHSDLALLRVPDDGFWEG VEAIRMEPEDVKLGELVTVI- GYPRGGEK ICLTKGIVS- RLHFNGEYLSIQIDAVRVPG RGERERGRE |
| AFGL01021447.1 | Full length | Y | XP_005852657.1 | MVRTFMAQPTREDAQVL- LAKRFFLMGC LGLPWL- WAVSVMYFWPRVYSGRSSSIG TGSTISSGVARRDGGNPELRK- WVGRSLV GAVVGFGLLFG- WVISFQLRWREWGSGV LNYMISAPESYSSGW |
| AFGL01012239.1 | Partial | N | WP_145058514.1 | MLLLLTLPHAVISNADAFH- LYPTIIIYRPQ QQQPT- NCLTSCSGRLFPSMSTP- SPPSR SLLAALLPLLLL- LLPFVTHGFVFPQPKLG NILPLPRVPSTPMASST- SPSSSSSSSASS SSSSSRASRT- SASPTFFRRRWRPLFQQQ PPAGEDPSGAFIPTDMP- PIPPSSPSPLSP NYKTI- AALIGGELLLGLLAFPLGHF- FKINP LATAKAAMGFG- PLLAVSALLTLPALLLHFA SGSEIEDMLRESLEGIFGS- DTAKKQPLVVI AISVLL- SLSAGIGEEVAFRGALQPIL- SQVT NNPWMGLGISS- FIFAALHAATPRYFLLAL LLSVYLGVLQLRYANLWLP- MMVHTLFDVV GFVQN- HFWGGGKGGSKGKTA |

Continued on next page

Table 5 continued from previous page

| Accession | Status | Contig end | Closest Homolog | Protease Sequence |
|---|---|---|---|---|
| AFGL01010117.1 | Partial | N | XP_005819270.1 | MASAPLSRGGFRRWL- GHGGLLVVIVLSS STSSS- LAFGPPLRTTLPPLH- LVSLPCGSS SVGSSSSQRISSP- CAPLSSPTPGVVKRR MLP- PLLTRPGSSEDPEALEVDLS- GQPRW RRATTRLITAWR- RRQQQRAFTTQSLQSS ASAMASPPGESGEAW- GASELPSYKIRPD WA- PAWVPTWAVTLHPIIQVVIM- VALYFFH MLVLSKVAMPF- PFQLVPNSHGLFQSIGM DSLAGLLVLGVMAYFRRSR- GLPLRPVVFG DRPLPFNPP- KKGRGKVWRILALLMVAYL TSGSASKWVEDVFGLSEL- NVALGRAVQV LAGH- LVWVFLGVAILKTQAGGGGR- GAFF GKKSVWMKSSFN- TYWLWWAIGGYYVSS LAFHVADTVNQVVL- PASAFEVDSVVTQM INPEG- NDVMAMLVGSLAPCVSAP- WWEE VLYRGFLLPALYLFYP- PDVATLVSAVIFAA HHMSMSG- MIPLTTLGYLWAVLYLLSGNL LVPIAIHSLWNSRVFLGSLLGI |
| AFGL01002094.1 | Partial | Y | XP_008861991.1 | MSADKLLFNYHD- DAQVTEGDQVLFKPP GWL- NDRCINFYFRVLEHEEFPDR- PDLLFM DPAVVSCMLIQCT- PRVTDQDELEELGEG LGL- DRRSMVFIPVNDAEEFFS- GCTHWYVR GRSLFLFL- HNHMANQSLHPPFGLVSFPKS TFLSLSLSLSPFLSLSLSRSH- SLFFLSPSCSL LLPPIYLLVLF- SLPRSSDSPSIQPRRPPPPAA AAAAAA |

Continued on next page

Table 5 continued from previous page

| Accession | Status | Contig end | Closest Homolog | Protease Sequence |
|---|---|---|---|---|
| AFGL01008528.1 | Partial | Y | XP_001022365.1 | MKLLGLSVAAALGLLAHT-GAALVRGSSGKASRL-SLDLEPHHHYTFER-YVQEFGKNYVRGSEE-WAERKALFLERL-RDIQAHNSAHSDGKK-TYRKGVNEYADWTEAEL-RGMLGFKPSGEWHRKRQP-GLETLSALMEGAGNESQLS-GFRQQERLLTNLPTHVD-WREKGAVTKPKNQGACGSS-CWTFSGAEAIESALWLST-GKLFTLSEQEFVSCTPN-PDHCGGTGGCYGATQDI-LYDYVLAKGFTLEKDYPY-ISGNGTEPVCLRQPLHVGSI-AGFVDVTPNDLGAHMLAIN-SQPLAISLDASEFHNYHS-GVLTFEVRREGGRE |
| AFGL01011939.1 | Partial | Y | XP_067800298.1 | MRRTDTISHNPHTTHT-TQLQPQPAMPAAGAADQP-PDGPSSSPPASSSSSSSP-PAPQPGTSKYAIRLFGPKD-DHSICKIYVEKAKKSPRVQELL-GAITDLGCPIDLDT-MVDCRKCNIAATGGFI-ANHQGLKEGDYKP-KVVVCEDNLLGSHYVAQT-LAHELVHAYDQCRAKVN-WRSCYHYACTEIRASTLS-GECDFIEEFDRGIFGM-RAQHQECVRRRAELSM-MVNESCKDRAKEVIEKVFAR-CYQDTAPYDKIP |
| AFGL01004592.1 | Partial | Y | XP_005855021.1 | MDCLKCKASLKAV-MARPRPKALLDALAET-GIQTLENAPESRR AALAASTQRHFQEGS GGGGGGKRGEGGSK ARGGAAAAAAGAGAG TTLNIKCLNCAEDGIA MKARAYMEAPPMSVVL CANRLQDRREMEEALV HELVHVYDVRT |

Continued on next page

Table 5 continued from previous page

| Accession | Status | Contig end | Closest Homolog | Protease Sequence |
|---|---|---|---|---|
| AFGL01013087.1 | Partial | Y | WP_298615612.1 | LTVAEVEKGSPAEAGG LIAGDKLLNVDGQAT TGLLIDDCANLLRGP VGTKVGVVAKSKTG ENKDLI-ITRQLIKIQ SVEASTATKG-GKKV GVIRIRSFSQT-TADT VAAELEKLEKGGIS ALVIDLRGNPGGFL GGGID-TARLFLPSYE KIVSVVGKT-GIVEEY QTTDEGRDTTK-PLF LVVDKRTASASEVLS AALKENGRAKLVGEK TFGKGVVQTIEQVGQ GAGVSVTIAAYSTPLG NNINKKGIEVDIPSAC EATD-MAVSCLPEGVI |
| AFGL01010546.1 | Partial | Y | XP_005856080.1 | SSVPPSSSSFSSLPP PMP-FISKETFPVIKPR VP-TQDNSCDCGVFVL KYGDMMLTDVLPTL GKWTQGMLNRKLRD VIKNDAFSQEDVRV MREELEEIFRRLERE YARGGFTEMEQKGG GGGGKAGGKKKEGG RTREEDEEREEEAEA ARVALALVEEMKAKE RREEEEKKKKKKEEK EEEEGEEGEEGEKR KEAEEEEEDDDATQV EEEEQQGRKEAGAG GREEGGGGKG |

Continued on next page

Table 5 continued from previous page

| Accession | Status | Contig end | Closest Homolog | Protease Sequence |
|---|---|---|---|---|
| AFGL01014613.1 | Full length | N | XP_008896704.1 | MMRSMARRICLA RPV-PLLQRQHSLA CRRPLSH-TALCDS RAPALNQVI-GARR AFSVETGGSAVT TEPPPPPPLDRG AGTTP-TATVAAD AVPPTAAT-PEVVA AALKPKELVEYLD RYIVGQADAKRAV AIAL-RNRWRRHK LPEDLRTEI-IPKNI LMIGPTGCGK-TEI ARRIAKLSDAPFIK VEATKFTEVGFHG RDVDQI-IRDLVEIS VNMTKKRRSEAIK AEAQKAVEEKILQI LT-GPYAREDAKETF RDLL-RKGSLNDRL IDVEV-PADRGGAG GVFDKA-GAAPGSG AFAEGNPNVSMA DFIGKFSRAIMPK RPMEKRK-ILVKEA MPLLLEVEMEKLV DMSDVTREAIQSV EES-GIVFIDEIDKIC NPSDYR-GADASAE GVQRDLLPLIEGCT ISTKHGNVNTDYILF ISS-GAFHSCKPADLL AELQ-GRLPIRVELKG LTEED-MYRILTEPVT NLIAQQVALL-STEKV ALEFTDEAVRE-VARV AALVNKTVENIGARR LHTVLERIVEEISF EASD-KPEGSVITVD KALVQS-RVNDLLVE SDISKYIL |
| AFGL01014773.1 | Partial | Y | XP_005856080.1 | LGSKPCLVFMDSLSVHRTDKI-HTALLHYLKQEWKKRKA |
| AFGL01015019.1 | Partial | Y | XP_002295819.1 | FREGITVVSAFEGYAFDAGM-RVGDSLLAVDDVPVA-GRSVDQVRDLLRGPPN-TQVKVKITRAGLPPPSQQEL-LLTRRLIRVPDVKLTSFLGPP-KDGIGHM |
| AFGL01017517.1 | Partial | Y | XP_005855041.1 | LLLVSGGGAAYEYGLASAQLE-QRLRRAGVRLTVCVD-KVAASGGYMMAVVADTLI-AAPFATVGSIGATTSVPVL-NVGGFLKRVGIQAYTVKS-GKYYFTFLPSLPPFLRLFNPL-FRLSSVLFLVLSNPSLPPSLL |

Continued on next page

Table 5 continued from previous page

| Accession | Status | Contig end | Closest Homolog | Protease Sequence |
|---|---|---|---|---|
| AFGL01009172.1 | Partial | Y | XP_005853719.1 | HNRIVYLGMALV-PAVTELIIAELLYLNYESQ-DRPIYMYINSPGIST-PDGRAMGFDTEAFAIADVM-SYIRAPIATVCVGQCYGSAAM-LLACGEKGQRYSLP-NGSIMLHQPRSLARGQAS-DIAIKAREVMRNRK-INCEMLAKACGK-PLDVIIKDASRKRYF-SPEEACEYGLIDQVLEST-NDLPMGLIPRKQGGDE-PENDLNMADVRGYS-NPAGYGDSADGNMPTTPF |
| AFGL01017519.1 | Partial | Y | XP_005855041.1 | PAFKRRGVTPAR-RIYRVGLKQSLVAR-LVMKATKAFNLTSSSSSS-NTTTTG-NITTTTSPFRALRSLPPHK-LASLSLKILLPLLLLYSLRHLIT-FKATMQTLMIISILAEIAGR |

[1] NCBI Nucleotide
[2] Status: Partial/Full Length.
[3] End of Contig (within 1kb)

### 2 Supplementary figures

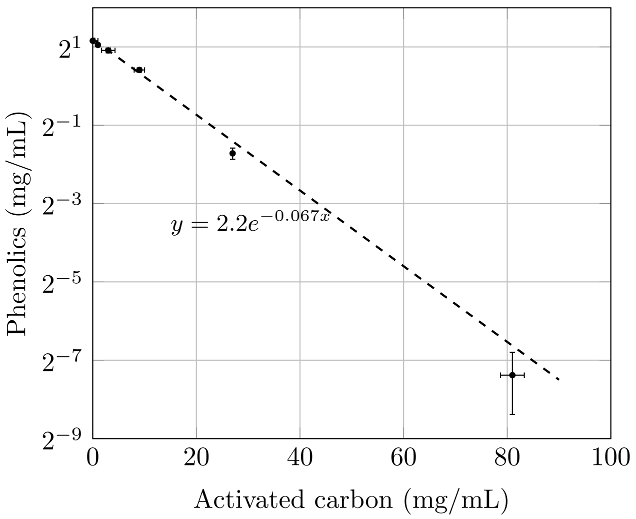


Supplementary figure 1: Phenolics removal profile from dPDF at increasing concentrations of activated charcoal.

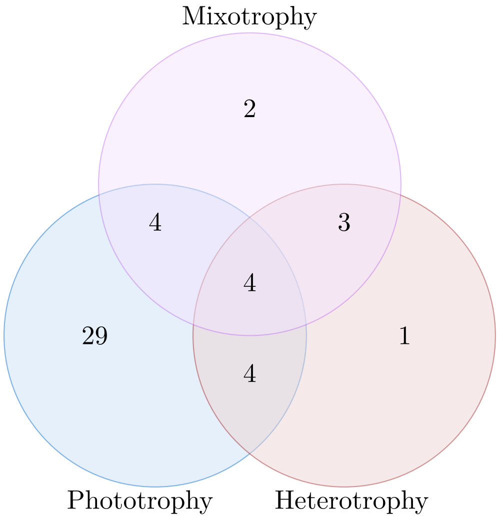


Supplementary figure 2: Venn diagram of proteins detected in the extracellular medium in trophic condition.
